# Biophysical and proteomic analyses suggest functions of *Pseudomonas syringae* pv *tomato* DC3000 extracellular vesicles in bacterial growth during plant infection

**DOI:** 10.1101/2021.02.08.430144

**Authors:** Martin Janda, Christina Ludwig, Katarzyna Rybak, Chen Meng, Egidio Stigliano, Leon Botzenhardt, Beata Szulc, Jan Sklenar, Frank L.H. Menke, Jacob G. Malone, Andreas Brachmann, Andreas Klingl, Silke Robatzek

## Abstract

Vesiculation is a process employed by Gram-negative bacteria to release extracellular vesicles (EVs) into the environment. Bacterial EVs contain molecular cargo from the donor bacterium and play important roles in bacterial survival and growth. Here, we describe EV production in plant-pathogenic *Pseudomonas syringae* pv. *tomato* DC3000 (*Pto* DC3000), the causal agent of bacterial speck disease. Cultured *Pto* DC3000 exhibited EV structures both on the cell surface and in the vicinity of bacterial cells, observed as outer membrane vesicle (OMV) release. We used in-solution trypsin digestion coupled to mass spectrometry to identify 369 proteins enriched in EVs recovered from cultured *Pto* DC3000. The predicted localization profile of EV proteins supports the production of EVs also in the form of outer-inner-membrane vesicles (OIMVs). EV production varied slightly between bacterial lifestyles and also occurred *in planta*. The potential contribution of EVs to *Pto* DC3000 plant infection was assessed using plant treatments and bioinformatic analysis of the EV-enriched proteins. While these results identify immunogenic activities of the EVs, they also point at roles for EVs in bacterial defences and nutrient acquisition by *Pto* DC3000.

## Introduction

Successful colonization of hosts depends on the ability of microbes to defend themselves against host immune responses and acquire nutrients. Bacterial pathogens use macromolecular translocation systems and deliver virulence proteins, so-called effectors, to circumvent host immunity (Buttner & Bonas, 2010). *Pseudomonas syringae* pv. *tomato* (*Pto*) DC3000 is the causal agent of bacterial speck, a common disease that occurs in tomato production worldwide (Mansfield *et al.*, 2012, Wilson *et al.*, 2002). *Pto* DC3000 is a Gram-negative bacterium that invades through openings in the plant surface and propagates in the apoplast, where it takes up nutrients and proliferates (Melotto *et al.*, 2006, Xin & He, 2013, Xin *et al.*, 2018). Plants respond rapidly to colonization by microbes, activating innate defence strategies, which can broadly be categorized into pattern-triggered immunity (PTI) activated by microbe-associated molecular patterns (MAMPs) and effector-triggered immunity (ETI) induced upon recognition of virulence factors or their actions (Couto D. & Zipfel, 2016, Dodds & Rathjen, 2010). Virulence of *Pto* DC3000 largely depends on the Type-III secretion system and its secreted effectors (Kvitko *et al.*, 2009, Nobori *et al.*, 2018, Nobori *et al.*, 2019, Nomura *et al.*, 2006). A number of Type-III secreted effectors from *Pto* DC3000 and their *in planta* targets have been identified, many involved in immune suppression and some with roles in gaining access to nutrients (Wei *et al.*, 2018, Xin *et al.*, 2018). For example, genes encoding proteins for siderophore biosynthesis are upregulated *in planta* (Nobori *et al.*, 2018). In addition, pathogenic bacteria need to adapt to the host environment, resisting the defences induced by the host immune system.

The survival of infectious Gram-negative bacteria is greatly enhanced by releasing extracellular vesicles (EVs), a process widely studied in the context of bacteria pathogenic to humans (Schwechheimer & Kuehn, 2015). During infection, bacterial EVs can counteract the effect of antimicrobial peptides (Roszkowiak *et al.*, 2019). They also perform immunomodulatory functions by delivering virulence factors to recipient cells resulting in immune-suppression (Kaparakis-Liaskos & Ferrero, 2015), despite having the capacity to activate defences due to their immunogenic cargoes (Kaparakis-Liaskos & Ferrero, 2015). More recently, a number of studies provide evidence that plant pathogenic bacteria, including cultured *Pto* bacteria release EVs (Bahar *et al.*, 2016, Chowdhury & Jagannadham, 2013, McMillan *et al.*, 2020). Although insights into both immunogenic and virulent roles have been achieved, little is currently known about the importance of EV production in bacterial infection success.

EVs are cytosol-containing membrane “nano” spheres that provide selection, storage and protection against degradation of enclosed cargoes in a highly dynamic and environmental cue-responsive manner (Bielska *et al.*, 2019, Rybak & Robatzek, 2019, Schwechheimer & Kuehn, 2015). Gram-negative bacteria actively form EVs by budding and shedding of the outer membrane, producing so-called outer membrane vesicles (OMVs) (Raposo & Stoorvogel, 2013, Roier *et al.*, 2016). Outer-inner-membrane vesicles (OIMVs) have also been described, involving a different mode of release such as endolysin-triggered cell lysis (Perez-Cruz *et al.*, 2015, Toyofuku *et al.*, 2019). EVs can also be produced in the form of elongated, tube-shaped vesicles as observed in Gram-negative *Francisella* spp. (McCaig *et al.*, 2013, Sampath *et al.*, 2018). As insufficient biomarkers are available to convincingly probe their origin, in particular for *P. syringae*, we will collectively refer to these vesicles as EVs. Notably, EV formation seems to be an essential process since no bacterial mutant lacking vesicle release has been reported so far and genetic reduction of vesiculation results in mutants with growth defects (McBroom *et al.*, 2006).

Previous studies revealed a number of molecular cargoes present in EVs from phytopathogenic bacteria of the *Agrobacterium tumefaciens, P. syringae, Xanthomonas campestris* and *Xylella fastidiosa* species. These EV-associated proteins include degradative enzymes, Type-II-secreted virulence-associated proteins, components of the Type-III secretion system and its secreted proteins (Feitosa-Junior *et al.*, 2019, Chowdhury & Jagannadham, 2013, Knoke *et al.*, 2020, Nascimento *et al.*, 2016, Sidhu *et al.*, 2008, Sole *et al.*, 2015). While respective genetic deletions of EV-associated degradative enzymes reduced bacterial virulence, the role of EVs in their delivery remained unanswered (Nascimento *et al.*, 2016, Sidhu *et al.*, 2008, Sole *et al.*, 2015). A previous seminal study described the production of EVs as a mechanism, by which *X. fastidiosa* regulates its attachment to host cells and thus the promotes systemic infection (Ionescu *et al.*, 2014). However, elongation factor Tu (EF-Tu) and lipopolysaccharides (LPS) are abundant components of EVs from *P. syringae, X. campestris, X. oryzae* and *X. fastidiosa* (Bahar *et al.*, 2016, Feitosa-Junior *et al.*, 2019, Chowdhury & Jagannadham, 2013, Sidhu *et al.*, 2008). Both represent MAMPs, with EV-associated EF-Tu shown to activate a prototypic PTI response in a receptor-dependent manner (Bahar *et al.*, 2016). Further results indicate that EV-induced immunity, triggered with EVs recovered from pathogenic *Pto* DC3000 and the commensal *P. fluorescens*, protects plants against *Pto* DC3000 infection (McMillan *et al.*, 2020). These studies hint at some contrasting roles that EVs from bacterial phytopathogens could play during plant infection.

Here, we used nanoparticle tracking analysis (NTA) to describe the production and the biophysical properties of EVs from *Pto* DC3000 in different growth conditions including their accumulation *in planta.* Analysis of *Pto* DC3000 cellular, outer membrane (OM) and EV proteomes by mass spectrometry identified 369 EV-enriched proteins. The potential contribution to bacterial growth *in planta* of these proteins was assessed using bioinformatic analysis as well as exploring plant responses to EVs. These findings expand our understanding of the functions of EVs in bacterial infection of plants.

## Results

### *Pto* DC3000 bacteria vesiculate and produce EVs in culture

We first examined the morphology of *Pto* DC3000 cultures by scanning electron microscopy (SEM). The bacteria displayed multiple spherical structures protruding from their cell surfaces, with diameters in the range of 20-120 nm (Fig. 1A; S1A). These vesicle-like structures appeared to be released from the surface, as similarly sized vesicular structures could also be observed in the vicinity of the bacteria (Fig. S1A). To determine whether these structures were released from the bacterial cell surface, supernatants of planktonic *Pto* DC3000 cultures were filtered through 0.22μm-pore membranes to remove intact bacteria and measured by Nanoparticle Tracking Analysis (NTA) before (fluid sample) and after sucrose density gradient centrifugation followed by ultracentrifugation (gradient-enriched sample) (Fig. S2A). Density gradient centrifugation is used to separate EVs from other extracellular materials (Klimentova & Stulik, 2015). NTA measures particle number (concentration), particle size (median diameter and distribution), and particle surface charge (mean ζ-potential). Both sample types exhibited a polydisperse sized population of spherical structures with a diameter ranging from ~ 50 to 200 nm and median sizes of 100 nm and 115 nm for fluid samples and gradient enriched samples, respectively (Fig. 1C, 1G, S1B). This could suggest that *Pto* DC3000 releases vesicles from different biogenesis routes.

**Figure 1.**
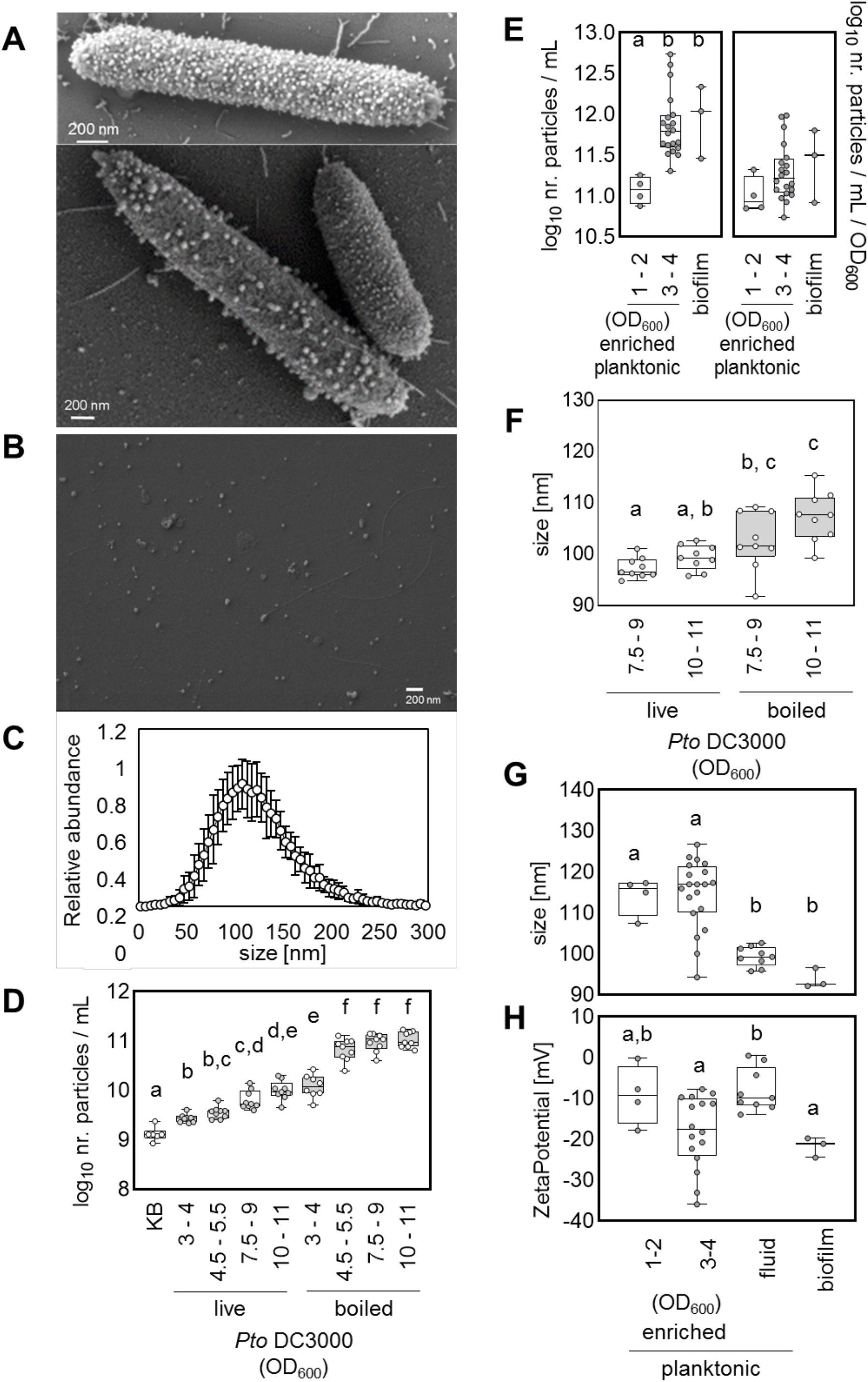
*Pto* DC3000 release extracellular vesicles with differing biophysical parameters depending on lifestyle. **A)** Representative SEM micrographs of *Pto* DC3000 growth in planktonic culture (OD_600_ = 3-4). **B)** Representative SEM micrograph of gradient enriched *Pto* DC3000 EVs purified from planktonic culture (OD_600_ = 3-4); scale bars represent 200 nm. For A and B, the micrographs were selected as representatives from three independent samples. For D-H each dot in the boxplot represents an independent sample. **C)** Size profile of gradient enriched EVs from *Pto* DC3000 planktonic cultures (OD_600_ = 3-4), the values represent mean and standard deviations from n = 20. Concentration **(D)** and size **(F)** of EVs from fluid samples before (live) and after heat inactivation of bacteria (boiled); King’s B (KB) medium. Concentration **(E)**, size **(G)** and ζ-potential **(H)** of EVs from planktonic cultures (enriched = gradient enriched OD_600_ = 1-2 and 3-4 and fluid samples OD_600_ = 7.5 - 11) and of EVs from biofilm cultures. For D and F n = 8-9 independent samples, for samples in E and G n = 4 for OD_600_ = 1-2 and n = 20 for OD_600_ = 3-4 for gradient enriched samples, n = 9 for fluid samples and n=3 for biofilm. The box in boxplots extends from 25^th^ to 75^th^ percentiles, whiskers go down to the minimal value and up to the maximal value, the line in the middle of the box is plotted at the median. Different letters indicate significant (Welsch’s ANOVA with Dunnett’s T3 multiple comparisons post hoc test; p < 0.05).

Pellets obtained from gradient enriched samples were further examined by SEM and revealed numerous spherical structures (Fig. 1B), yet in this analysis the vesicles diameter ranged between 25 and 170 nm with a median around 50 nm. It is possible that conditions used for SEM and NTA differ in their capacity to hydrate the vesicles and/or that NTA underestimates smaller particles (Bachurski *et al.*, 2019). In addition, co-purifying filamentous structures could be detected (Fig. 1B). To determine whether EV production is an active process, EVs were quantified from culture supernatants of *Pto* DC3000 over cultivation time, with increasing particle numbers observed with bacterial density (Fig. 1D, 1E, S2B). While the total amount of EVs recovered from bacteria at late exponential growth was higher compared with early growth stages (Fig. 1D, 1E), calculation of the amount of EVs produced per bacteria showed that numbers were similar between growth stages (Fig. 1E). The median diameter and ζ-potential of EVs was comparable across growth stages (Fig. 1F, 1G, 1H).

Quantification of EVs from *Pto* DC3000 cultures that were incubated in fresh media followed by heat inactivation showed increased vesicle numbers (Fig. 1D). This suggests that heat inactivation could additionally trigger the production of vesicles, i.e. from cellular debris and/or through a process described as explosive cell lysis (Toyofuku *et al.*, 2019). Given this increase and the distinct size of the vesicles from heat inactivated bacterial cultures (Fig. 1F), it suggests that the vesicles recovered from culture samples without heat inactivation are predominantly produced from bacteria as an active process.

EVs were also isolated from biofilm grown *Pto* DC3000 cultures (Fig. S2C, S2D). The median diameter of these EVs was smaller compared with EVs from planktonic gradient enriched *Pto* DC3000 EVs but had a similar size to EVs from fluid samples (Fig. 1G). The mean ζ-potential of EVs from biofilm samples was similar to EVs from planktonic gradient enriched samples but more negative than EVs from fluid samples of planktonic *Pto* DC3000 cultures (Fig. 1H). Thus, EVs show diverse biophysical properties depending on bacterial lifestyle, further refuting the possibility that the purified particles are solely formed through nonspecific assembly of shed membrane fragments.

To investigate whether the biophysical parameters of *Pto* DC3000 EVs could change upon mechanical treatments, we subjected the samples to sonication, heating and freezing. In addition, we tested the effect of incubation with Proteinase K, a treatment used to deplete the EV samples of extravesicular proteins (Metruccio *et al.*, 2016). None of the treatments significantly affected the particle concentration (Fig. 2). Also, particle size was not significantly changed upon sonication and Proteinase K treatments (Fig. 2A, 2D). A significant increase in particle size was observed after ten freeze-thaw cycles and longer heat exposure (Fig. 2B, 2C). These observations suggest that EVs are affected by more extreme temperature treatments, maybe forming higher aggregates, while sonication, shorter heat incubation, fewer freeze-thaw cycles and Proteinase K treatments showed no significant effects.

**Figure 2.**
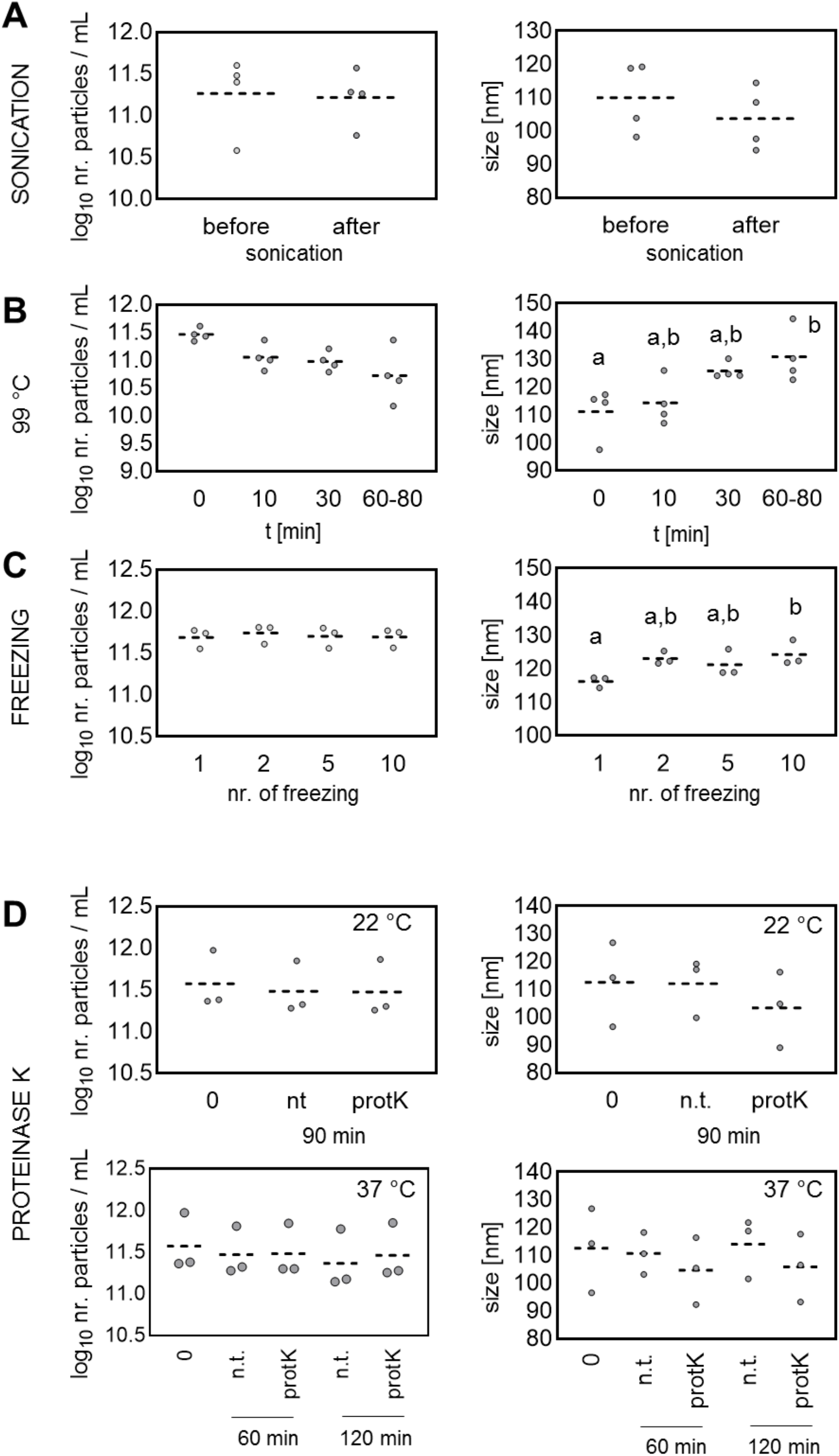
Disruptive treatments cause minor changes to biophysical parameters of gradient enriched EVs. Concentration and size analysis of *Pto* DC3000 EVs from planktonic cultures (OD_600_ = 3-4). Effects of **A)** sonication (10 times 30 s); **B)** 99 °C for 80 min; **C)** freezing and thawing (up to ten times); **D)** Proteinase K treatment (10 μg/mL) at 22 °C and 37 °C for up to 120 min. Individual circles represent particles characteristics from independent samples; n = 3 to 4. Different letters indicate significant difference (One-way ANOVA with Tukey post hoc test; p < 0.05).

### *Pto* DC3000 bacteria produce EVs *in planta*

To determine whether *Pto* DC3000 releases vesicles *in planta*, apoplastic fluids were recovered from *Pto* DC3000-infected and control-treated *A. thaliana* leaf tissues at different time points. The apoplastic fluids were collected, filtered to remove intact bacteria and then directly characterized by NTA without density gradient centrifugation and ultracentrifugation. In these apoplastic fluid samples, we identified particles with a median diameter of ~96 nm (Fig. 3A), which increased in abundance upon infection with *Pto* DC3000 (Fig. 3B), consistent with previous findings (Rutter & Innes, 2017). Increased particle abundance correlated with the bacterial infection time and titers (Fig. 3B, Fig. S3A, S3B). We also analysed EVs from apoplastic fluids of plants, which were co-treated with 100 nM flg22 and *Pto* DC3000. Particle numbers were lower than those recovered from *Pto* DC3000 infection only, consistent with induced plant defences (Fig. S3C). Taken together, the higher particle numbers and polydisperse particle size isolated from *Pto* DC3000-infected plants compared to flg22 immune-stimulated plants hints at bacterial-derived EVs present in *A. thaliana* apoplastic fluids (Fig. 3B).

**Figure 3.**
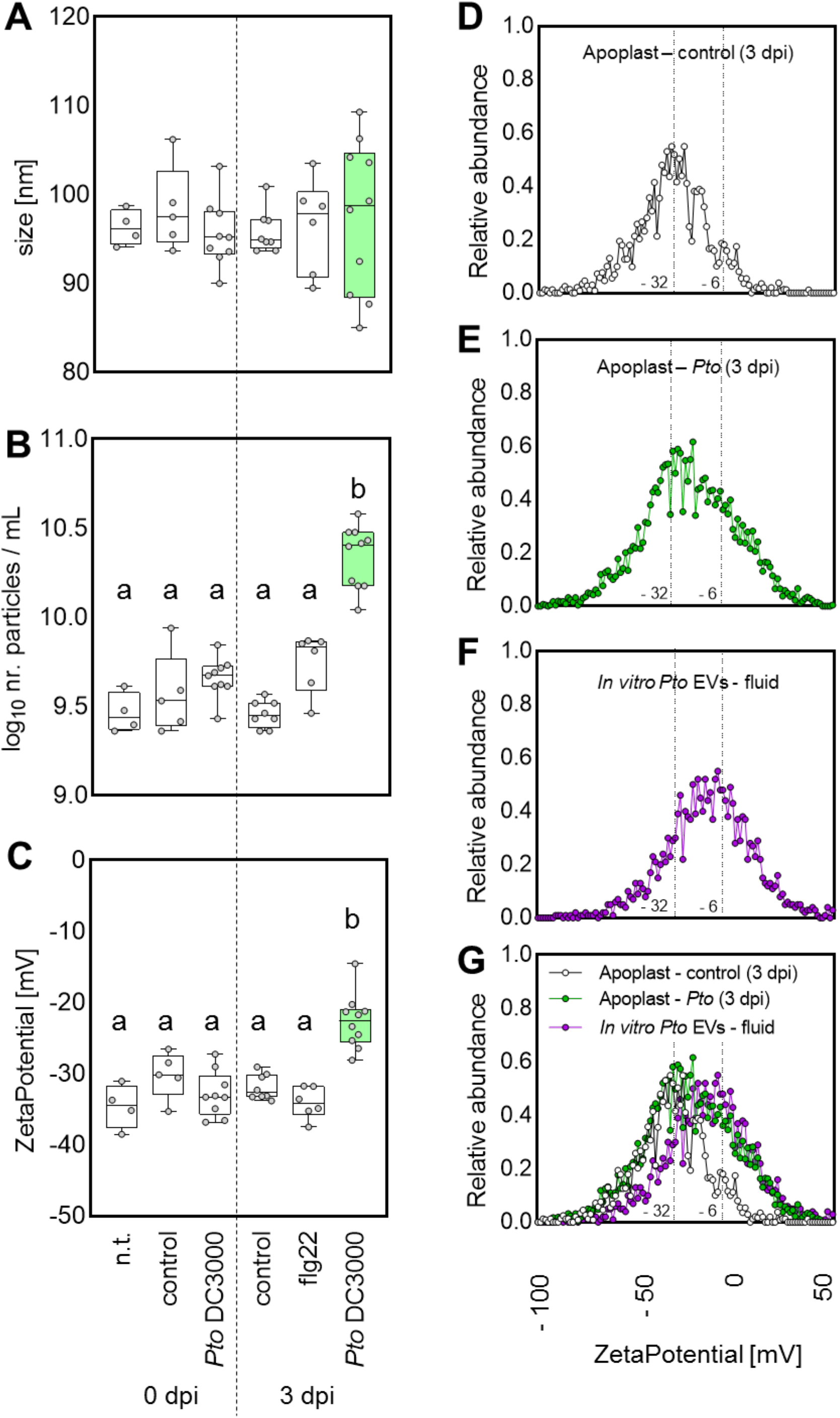
*Pto* DC3000 release EVs *in planta*. Size, concentration and charge measurements of apoplastic fluids from *A. thaliana* plants infected with *Pto* DC3000 and treated as indicated. **A)** Size of the particles. **B)** Particle concentration in apoplastic fluids. **C)** ζpotential of the particles. For A-C the variants represent: n.t. – non-treatment; control – 10 mM MgCl2; *Pto* DC3000 (OD_600_ = 0.0006); 100 nM flg22. **D-G)** The profile of ζ-potential for particles detected in Arabidopsis apoplast treated with MgCl2 (control; D) for 3 days, with *Pto* DC3000 (*Pto*; OD_600_ = 0.0006; E) for 3 days and for EVs from *Pto* DC3000 grown in culture (fluid; F). The dots represent the mean from 8 (D, G_white), 10 (E, G_green) and 13 (F, G_purple). For A-C each dot in the boxplot represents an independent sample. The box in boxplots extends from 25^th^ to 75^th^ percentiles, whiskers go down to the minimal value and up to the maximal value, the line in the middle of the box is plotted at the median. Different letters indicate significant difference (One-way ANOVA with Tukey post hoc test; p < 0.05). The green colour is highlighting the particles from *Pto* DC3000 infected plants (3 dpi).

Since *Pto* DC3000 (fluid sample) and *A. thaliana* (apoplastic fluid samples) EVs did not significantly differ in diameter (Fig. 1G, 3A), we focused on the charge of EVs, reflecting the different surface composition of bacterial (prokaryotic) and plant-derived (eukaryotic) EVs. Evaluation of the mean ζ-potential identified significantly less negatively charged EVs recovered from apoplastic fluids of *Pto* DC3000-infected plants at three days post infection compared with control treatments and earlier time points (Fig. 3C, 3D, 3E, 3G). This time point correlated with *in planta* bacterial proliferation and depended on bacterial inoculum (Fig. S3A, S3B). Plotting the relative particle abundance over particle charge, the ζ-potential profiles of EVs recovered from apoplastic fluids of untreated, control-treated and flg22-treated *A. thaliana* identified major peaks around −32 mV (Fig. 3C, 3D, S3J). By contrast, the ζ-potential profile of EVs recovered from apoplastic fluids of *Pto* DC3000-infected *A. thaliana* had a broader distribution with a similar major peak around −32 mV and an additional shoulder around −10 mV (Fig. 3E, 3G). Comparison of the different ζ-potential profiles revealed similarities of the major −32 mV peak across all plant samples, likely representing a plant-derived EV pool (Fig. 3C, 3D, S3J, S3K). Notably, the shoulder around −10 mV detected from apoplastic fluids of *Pto* DC3000-infected plant samples showed large overlay with the ζ-potential profile of EV recovered from *Pto* DC3000 cultures (fluid samples), with a peak from −20 to 0 mV (Fig. 3E, 3F, 3G). This could, therefore, represent a bacterial-derived EV pool. Since the ζ-potential profiles of EVs recovered from apoplastic fluids of flg22-treated *A. thaliana* did not differ between untreated or control-treated leaves (Fig. 3C, S3J, S3K), we found no evidence that plant EVs modulate their surface charge during infection. Thus, our data strongly suggest that *Pto* DC3000 release EVs during plant infection.

### EV samples purified from *Pto* DC3000 cultures trigger plant immune responses

Bacterial EVs contain immunogenic molecules such as EF-Tu and LPS, and can contain digestive enzymes and effectors that undermine host defences (Bahar *et al.*, 2016, Feitosa-Junior *et al.*, 2019, Knoke *et al.*, 2020, Rybak & Robatzek, 2019). To determine the effect of EVs from *Pto* DC3000 on plant cells, we first examined the ability of the *Pto* DC3000 EVs to modulate the outcome of bacterial infection. We pre-treated *A. thaliana* leaves with *Pto* DC3000 EVs, which limited the growth of subsequently infected *Pto* DC3000 bacteria *in planta* (Fig. 4A, S4A). Thus, the immunogenic potential of *Pto* DC3000 EVs is sufficient to restrict bacterial colonization, consistent with recent observations (McMillan *et al.*, 2020). Since MAMP treatment mediates anti-bacterial protection through the induction of plant immune reactions (Zipfel *et al.*, 2004), we next evaluated defence gene expression to EV treatment using *pFRK1::GUS* reporter lines (Kunze *et al.*, 2004). Seedlings were treated with purified EVs isolated from *Pto* DC3000 cultures, and GUS staining was measured after 18 h. We observed a significant induction of *pFRK1::GUS* expression triggered by the EVs albeit lower when compared to flg22 treatments (Fig. 4B, 4C). EV-induced *FRK1* upregulation is in agreement with previous observations (Bahar *et al.*, 2016). We also tested whether treatment with *Pto* DC3000 EVs could arrest seedling growth, a prototypic PTI response of plants to continual MAMP stimulation (Bredow *et al.*, 2019). Unexpectedly, we observed no significant growth reduction (Fig. 4D). This suggests that immune induction by *Pto* DC3000 EVs does not affect plant growth, unlike treatment with flg22 (Fig. S4B) (Bredow *et al.*, 2019).

A previous study demonstrated that bacterial (*X. campestris*) EV activation of *FRK1* expression depends on the EF-Tu Receptor (EFR), which is responsible for detection of the immunogenic peptide elf18 derived from bacterial EF-Tu (Bahar *et al.*, 2016, Zipfel *et al.*, 2006). To determine the pathway, by which the *Pto* DC3000 EVs trigger immune responses, we treated *efr-1* and *flagellin sensing 2* (*fls2*) mutants, the latter responsible for recognition of the immunogenic peptide flg22 of bacterial flagellin in *A. thaliana* (Zipfel *et al.*, 2004), with *Pto* DC3000 EVs and monitored *FRK1* gene expression. The *Pto* DC3000 EVs triggered *FRK1* gene expression in wild type and *efr-1* mutants to similar levels (Fig. 4E). No *FRK1* induction was observed in *fls2* mutants. Thus, the EVs isolated from *Pto* DC3000 cultures must contain bacterial flagellin. Notably, SEM analysis of gradient enriched EV samples showed the co-purification of filament-like structures (Fig. 1B), which could represent detached bacterial flagellar or pili. This suggests that co-purifying flagellin molecules may trigger plant immune responses.

**Figure 4.**
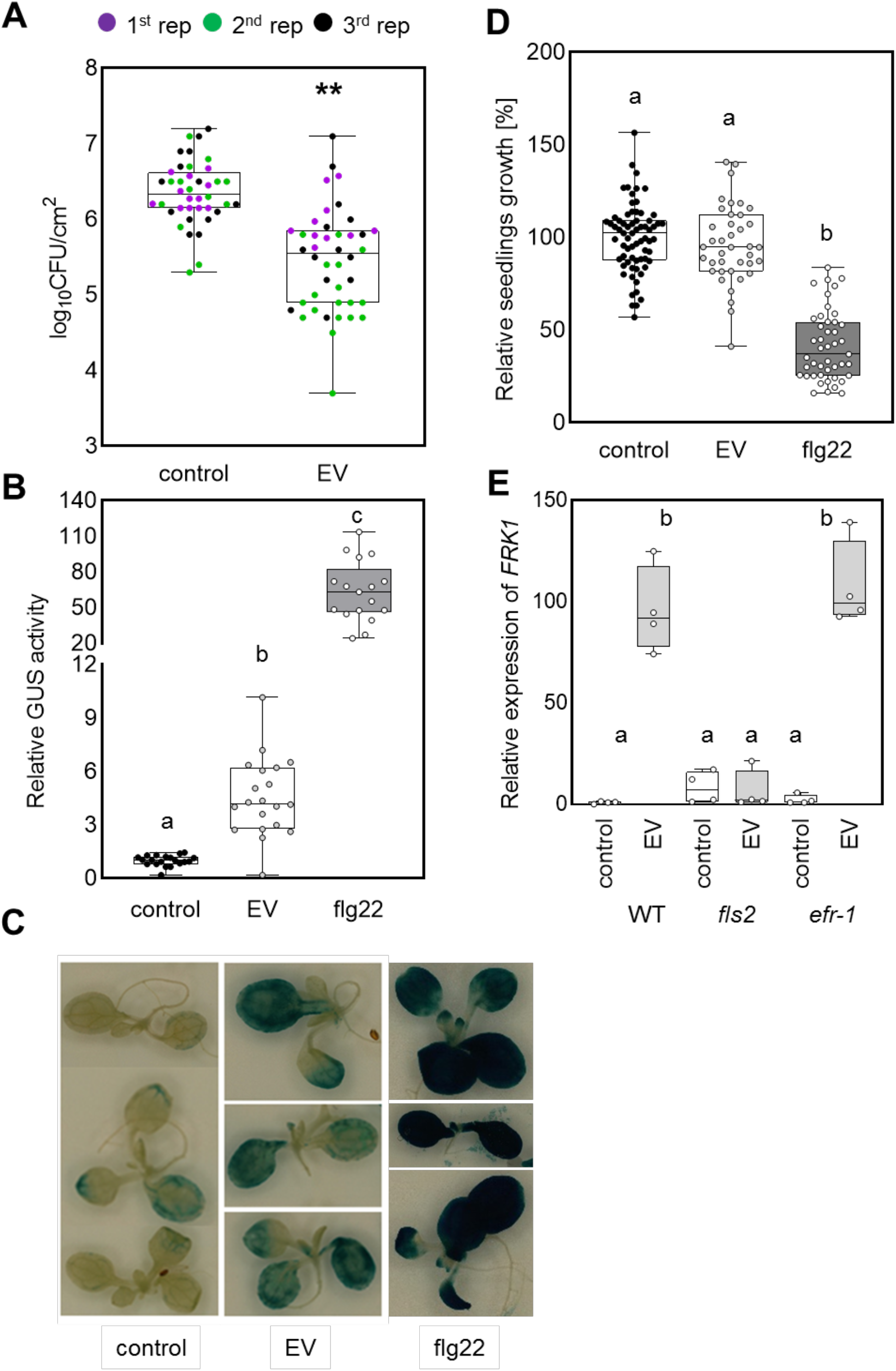
Immunogenic effects of *Pto* DC3000 EVs. **A)** *Pto* DC3000 growth (CFU) after infection into leaves of *A. thaliana* without and with EV pre-treatment at 3 dpi (control = 0.02 mM EDTA). Three biological repeats consisting each 12 independent samples were performed. The dots with the same colour represent independent samples from one biological repeat. **B)** Quantification of *pFRK1*::GUS activity in seedlings incubated without and with EVs (concentration ≈ 1.10^10^) or with 100 nM flg22 for 18 h. **C)** Representative pictures of *pFRK1*::GUS seedlings incubated without and with EVs (concentration ≈ 1.10^10^) or with 100 nM flg22 for 18 h. **D)** Fresh weight of seedlings grown without and with EVs (concentration ≈ 1.10^10^) for 8 days. For control n = 69; for 100 nM flg22 treatment n = 44 and for EV treatment n = 39 of independent samples. **E)** Relative *FRK1* gene expression in seedlings of the indicated genotypes incubated without and with EVs (concentration ≈ 1.10^10^) for 5 h (control = 0.02 mM EDTA), four independent samples were used for each variant. For A, B, D, E each dot in the boxplot represents an independent sample. The box in boxplots extends from 25th to 75th percentiles, whiskers go down to the minimal value and up to the maximal value, the line in the middle of the box is plotted at the median. Asterisks indicate statistical significances (two tailed Welsch’s t-test; p < 0.01) in A; different letters indicate significant differences (Welsch’s ANOVA with Dunnett’s T3 multiple comparisons post hoc test; p < 0.05)) in B, D and E.

### EVs from cultured *Pto* DC3000 are enriched in proteins with predicted roles in transport and antimicrobial peptide resistance

To gain insights into the function of *Pto* DC3000 EVs during the infection process, we characterized the proteome of EVs using liquid chromatography-based tandem mass spectrometry (LC-MS/MS). The *Pto* DC3000 EV-associated proteins were isolated from planktonic *Pto* DC3000 cultures by gradient enrichment. The proteomes of whole cells (WC) (Park *et al.*, 2014) and the outer membrane (OM) (Choi *et al.*, 2011) from bacteria grown to late exponential phase (OD_600_ = 3-4; Fig. S2B) were also analysed and compared with the EV proteome. As expected, we detected the most proteins from the WC sample (n = 1587), followed by the EV sample (n = 890) and 212 proteins in OM samples (Fig. 5A, Table S1). In total, 2898 proteins were identified over all samples, of which 1899 proteins were identified at least in three samples per sample type (WC, EV or OM). These proteins were taken forward for further analysis (Table S1). Similar protein intensity distributions were obtained for all samples (LFQ values were generated by MaxQuant, Fig. S5) and the four replicate measurements per sample type fell into sample clusters on the first and second principal components, suggesting a systematic difference in the proteomes of these three sample types (Fig. 5B). By comparing the proteomes of EV and WC, we identified 369 EV-enriched proteins, consisting of 162 proteins exclusively identified in at least three replicates of EV sample (EV unique; Fig. 5C) and 207 proteins significantly higher in the EV compared with WC (Fig. 5C; Table S1).

**Figure 5.**
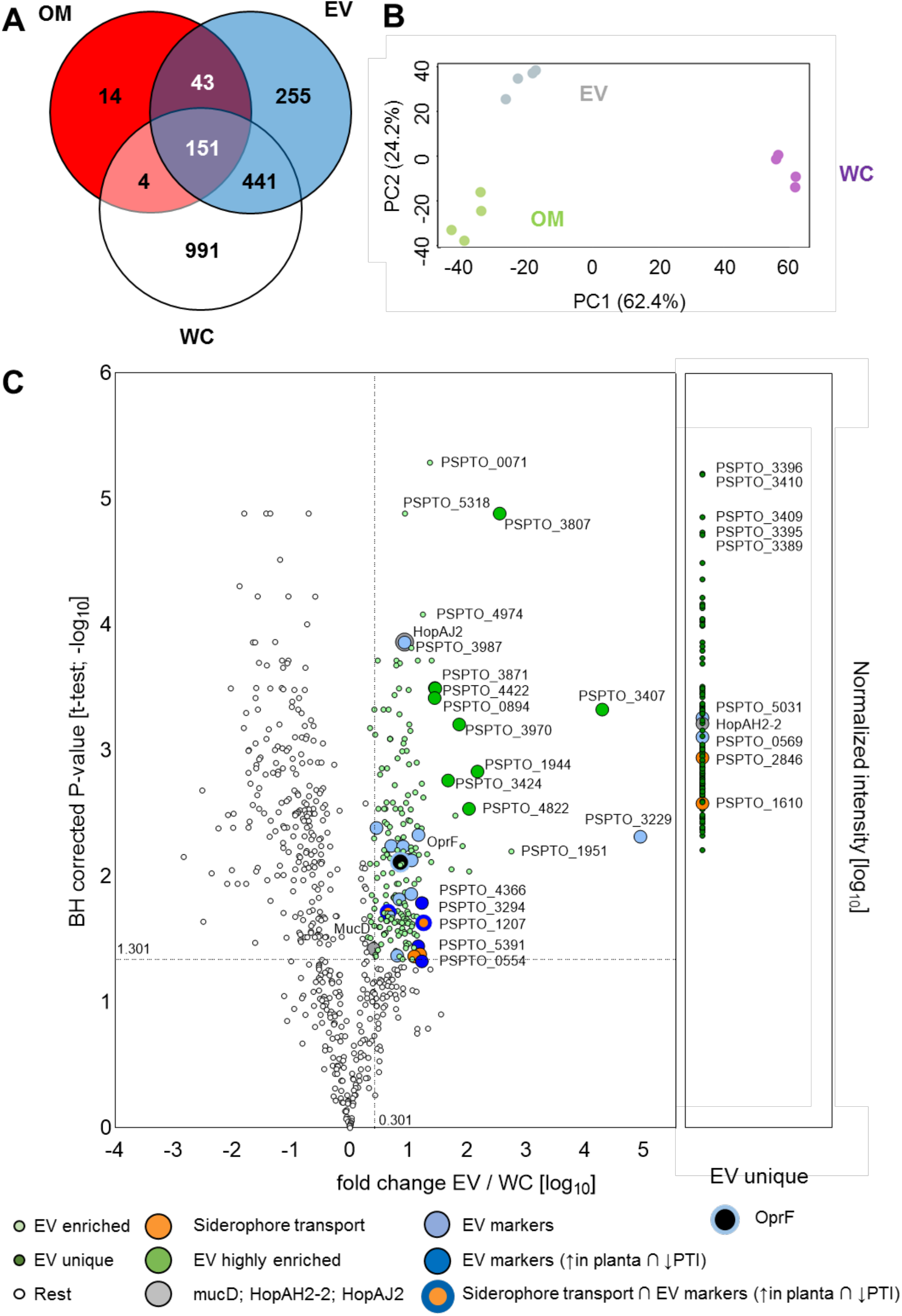
Proteomic analysis identifies 369 proteins enriched in *Pto* DC3000 EVs. **A)** Comparison of proteins detected in *Pto* DC3000 whole cell lysate (WC), outer membrane (OM) and extracellular vesicles (EV). **B)** Principal Component Analysis (PCA) analysis of identified proteins **C)** Volcano plot comparing EV and WC proteomes. EV-enriched proteins were defined in two categories: I) fold change EV/WC > 2 & FDR < 0.05 (t-test); II) measured in three replicates in EV but not in WC. In addition, the mean intensity in EV protein needs to be in the top 50% of all proteins, so only high intensity proteins in EV are selected. Four types of proteins enriched or unique in EVs were highlighted: proteins related to virulence; proteins related to siderophore transport; candidate EV biomarkers, and proteins highly enriched in EVs compared with WC.

Of the nine labelled highly-enriched EV and the five EV-unique proteins (Fig. 5C), eight have unknown subcellular localization (PSPTO_3807; PSPTO_4822; PSPTO_0894; PSPTO_3407; PSPTO_3424; PSPTO_3970, PSPTO_3396; and PSPTO_3409). This could indicate that these proteins are predominantly present in EVs, a localization not included in the predictions. Four proteins are annotated as lipoproteins, reported in other Gram-negative bacteria to mediate the cross-linking of the peptidoglycan (PG) layer with the OM and thus playing roles in the production of OMVs (Schwechheimer and Kuehn, 2015). PSPTO_3409 is the locus tag for the ATP-dependent ClpP-1 protease. Clp proteases were previously shown to regulate quorum sensing, in turn affecting OMV production and biofilm formation (Figaj *et al.*, 2019, Hall *et al.*, 2017). The other were annotated as hypothetical proteins.

The EV-enriched proteome included proteins related to virulence (Fig. 5C, Table S1), such as MucD (PSPTO_4221) (Wang *et al.*, 2019), HopAJ2 (PSPTO_4817) (Vinatzer *et al.*, 2006) and HopAH2-2 (PSPTO_3293) (Lovelace *et al.*, 2018, Schechter *et al.*, 2006). A major function of virulence proteins is the suppression of PTI (Block & Alfano, 2011). To test whether *Pto* DC3000 EVs could suppress a prototypic PTI response, we pre-treated leaves with EVs from cultured bacteria 24 h before eliciting a ROS burst with MAMPs (flg22, elf18). EV pre-treatments neither significantly reduced nor increased the MAMP-induced ROS production (Fig. 6A). This suggests that under the tested conditions, *Pto* DC3000 EVs are not predominantly involved in immune inhibition and/or further enhancing MAMP responses.

**Figure 6.**
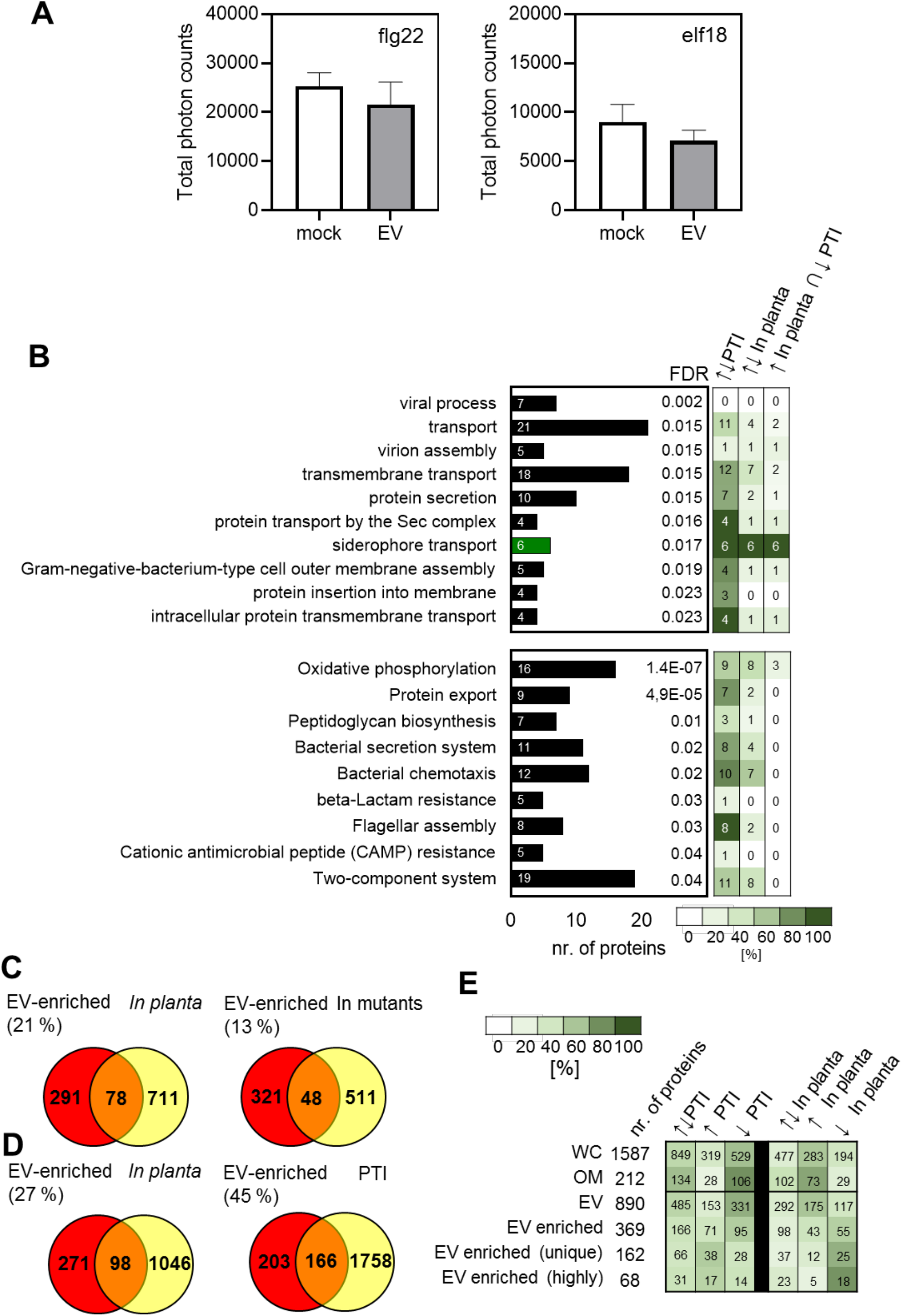
Protein profiles of EVs suggest functions other than classical immune suppression. **A)** Quantification of MAMP-induced ROS in leaves treated without and with EVs (concentration ≈ 1.10^10^) for 24 h. The bars represent mean and error bars represents SEM from n = 12. **B)** Enriched proteins in GO biological processes and KEGG categories in EV-enriched proteins. The heat-map represents the transcriptional profile based on the comparison of our data with transcriptomic data from (Nobori *et al.*, 2018) of genes encoding proteins identified by GO enrichment analysis. The intensity of green colour represents the percentage of affected genes and numbers in boxes represent the exact number of affected proteins. The arrows indicate transcriptional up- and downregulation. **C)** The Venn diagram represents the number of genes encoding EV-enriched proteins that are transcriptionally regulated *in planta* and responding to immune-deficient *in planta* conditions (Nobori et al. 2020). **D)** Transcriptional regulation of *Pto* DC3000 genes encoding EV-enriched proteins *in planta* and responding to PTI (Nobori *et al.*, 2018). **E)** The heat-map represents the transcriptional regulation of genes encoding identified proteins by proteome analysis *in planta* and responding to PTI (Nobori *et al.*, 2018).

Next, we performed a gene set analysis on the 369 EV-enriched proteins to examine the biological processes (from gene ontology (GO) (Ashburner *et al.*, 2000, The Gene Ontology, 2019) and pathways (KEGG (Kanehisa *et al.*, 2010)), in which these proteins are involved. In total ten GO biological processes were signficantly enriched (FDR < 0,05; DAVID bioinformatics resources (Huang da *et al.*, 2009a, Huang da *et al.*, 2009b)), out of which six were connected with the general process “transport”, including transmembrane transport, intracellular transmembrane transport, protein secretion, siderophore transport and protein transport by the Sec complex (Fig. 6B; Table S2). Nine KEGG categories, including cationic antimicrobial peptide resistance, β-lactam resistance and bacterial secretion system are significantly enriched in EVs (Fig. 6B, Table S2).

We used available *Pto* DC3000 proteome and transcriptome data to examine the *in planta* responses of the EV-enriched proteins (Nobori *et al.*, 2018, Nobori *et al.*, 2020). Comparison with the proteome data showed that of the 369 EV-enriched proteins 78 (21 %) are modulated *in planta* and 48 (13 %) are modulated in immune deficiency mutants (Fig. 6C, Table S3). Of the 369 genes coding for EV-enriched proteins, 98 genes (27 %) were differentially transcribed *in planta* compared to cultured bacteria and 166 genes (45 %) responded to the induction of PTI (Fig. 6D, Table S3). Most EV-unique and EV-highly enriched proteins were transcriptionally upregulated *in planta* whereas the majority of the genes of all identified proteins (WC, OM and EV) were downregulated *in planta* (Fig. 6E). When focussing on GO terms, we found that genes connected with the general process “transport” responded strongly to *in planta* conditions upon PTI activation (Fig. 6B). Interestingly, all genes connected with siderophore transport were strongly upregulated in response to *in planta* conditions, but downregulated *in planta* upon activation of PTI (Fig. 6B, Table 1). Thus, during sucessful infection the significant enrichment of siderophore transport proteins at EVs may suggest a role for EVs in iron or other metal ion acquisition (Fig. 5C, orange labelling). When focussing on KEGG pathways, genes connected with protein export (seven out of nine), secretion systems (eight out of eleven), chemotaxis (ten out of twelve) and flagellar assembly (eight out of eight) were affected *in planta* in response to PTI (Fig. 6B).

**Table 1.**
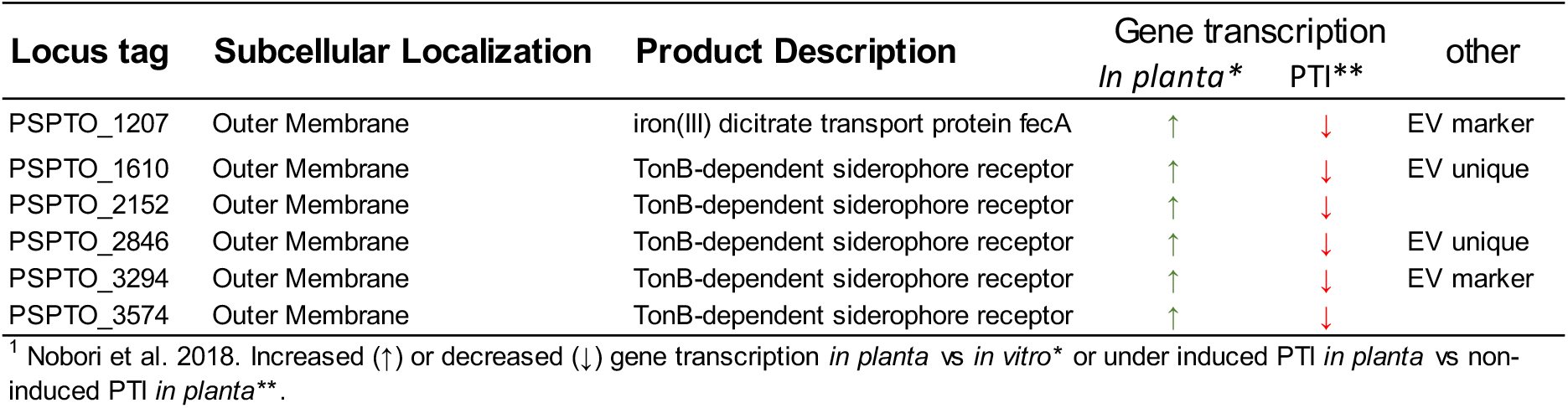
*In planta* transcription^1^ of genes coding proteins enriched in EVs belonging to GO:siderophore transport

| Locus tag | Subcellular Localization | Product Description | Gene transcription |  | other |
| --- | --- | --- | --- | --- | --- |
|  |  |  | *In planta* * | PTI** |  |
| PSPTO_1207 | Outer Membrane | iron(III) dicitrate transport protein fecA | ↑ | ↓ | EV marker |
| PSPTO_1610 | Outer Membrane | TonB-dependent siderophore receptor | ↑ | ↓ | EV unique |
| PSPTO_2152 | Outer Membrane | TonB-dependent siderophore receptor | ↑ | ↓ |  |
| PSPTO_2846 | Outer Membrane | TonB-dependent siderophore receptor | ↑ | ↓ | EV unique |
| PSPTO_3294 | Outer Membrane | TonB-dependent siderophore receptor | ↑ | ↓ | EV marker |
| PSPTO_3574 | Outer Membrane | TonB-dependent siderophore receptor | ↑ | ↓ |  |

<sup>1</sup> Nobori *et al.* 2018. Increased (↑) or decreased (↓) gene transcription *in planta* vs *in vitro*\* or under induced PTI *in planta* vs non-induced PTI *in planta*\*\*.

### *Pto* DC3000 appears to release EVs in the form of OMVs and OIMVs

Classification of the proteins enriched in EVs by putative subcellular localization revealed distinct localization profiles compared with WC and OM proteins. While 66 % of WC proteins were cytoplasmic, about half (51 %) of the EV-enriched proteins were cytoplasmic membrane-associated, with the next largest known class being OM-associated (11 %) (Fig. 7A). Yet, being putative localizations, we cannot exclude the possibility of other/additional localizations of the proteins as currently predicted, in particular for anchor-less proteins. Because the localization profile of the EV-enriched protein suggested that *Pto* DC3000 produces EVs in the form of OIMVs (Perez-Cruz *et al.*, 2015), we performed additional transmission electron microscopy (TEM) analysis of *Pto* DC3000 bacteria. Micrographs of the sectioned samples showed several structures reminiscent of budding vesicles from the bacterial outer membrane (Fig. 7B, 7C). Combining the data from proteomics and TEM, it may suggest that *Pto* DC3000 produces EVs in the form of both OMVs and OIMVs, as previously described for the closely related species *Pseudomonas aeruginosa* (Toyofuku *et al.*, 2019).

**Figure 7.**
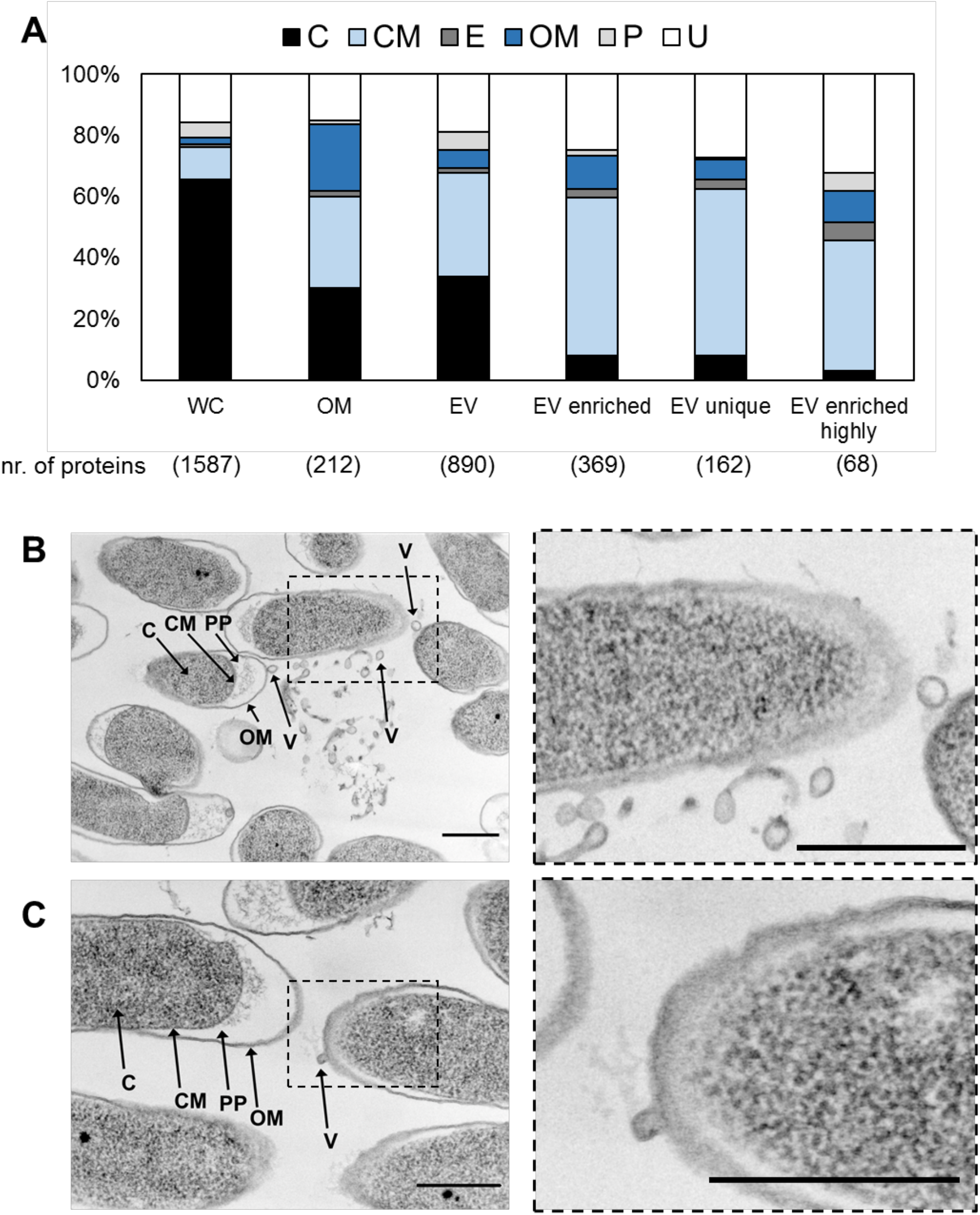
Localization profiles of EV proteins and TEM analysis suggest the release of *Pto* DC3000 EVs in the form of OMVs and OIMVs. **A)** Predicted protein localization of identified in whole cell lysate (WC), outer membrane (OM), EVs, EV-enriched, EV unique (the proteins identified only in EVs not in WC) and EV enriched – highly (protein which FDR < 0.005 and EV/WC > 20) in [%]. **B-C)** The left panel shows representative TEM micrographs from planktonic *Pto* DC3000 cultures (OD_600_ = 3-4). C, cytoplasm; CM, cytoplasmic membrane; OM, outer membrane; PP, periplasm; V, vesicle. All scale bars = 500 nm. **B)** A lot of smaller and larger vesicles in proximity to cells. It is important to note that the larger vesicle-like structures could also represent debris of death cells. **C)** Budding vesicle in the right part of the micrograph. Dashed boxes indicate enlarged regions of the micrographs shown in the right panel.

### Comparative analysis of proteomic data revealed 20 candidates for *Pto* DC3000 EVs markers

Since a number of EV proteomes have been reported from *P. aeruginosa* (Couto N. *et al.*, 2015, Choi *et al.*, 2011, Reales-Calderon *et al.*, 2015), we addressed whether the protein composition of EVs from *Pto* DC3000 and *P. aeruginosa* PAO1 (*Pa* PAO1) share similarities. We focussed on three published *Pa* PAO1 EV proteomes and found that 103 proteins were identified in the EV proteomes across the three reports (Couto N. *et al.*, 2015, Choi *et al.*, 2011, Reales-Calderon *et al.*, 2015). Of the 103 shared EV proteins from *Pa* PAO1, we could identify 100 orthologous proteins encoded in the *Pto* DC3000 genome and 44 proteins were enriched in *Pto* DC3000 EVs (Table S4). We refer to these as the EV “core”. These proteins were highly enriched in localization to the outer membrane (44 %) and cytoplasmic membrane (26 %) (Fig. 8A), consistent with EVs released in the form of OMVs (Fig. 7B, 7C). From these 44 proteins, 20 were putative outer membrane-localized proteins and thus represent good candidate biomarkers for the detection of EVs (Table 2, S4; Fig. 5C blue labelling). Interestingly, 20 out of 31 proteins with predicted membrane localization (20 out of 31) were transcriptionally regulated *in planta* in response to PTI activation (Nobori *et al.*, 2018), of which 14 showed downregulation. Overall, twelve of the 31 proteins responded transcriptionally to the *in planta* condition, with nine showing upregulation (Fig. 8B).

**Figure 8.**
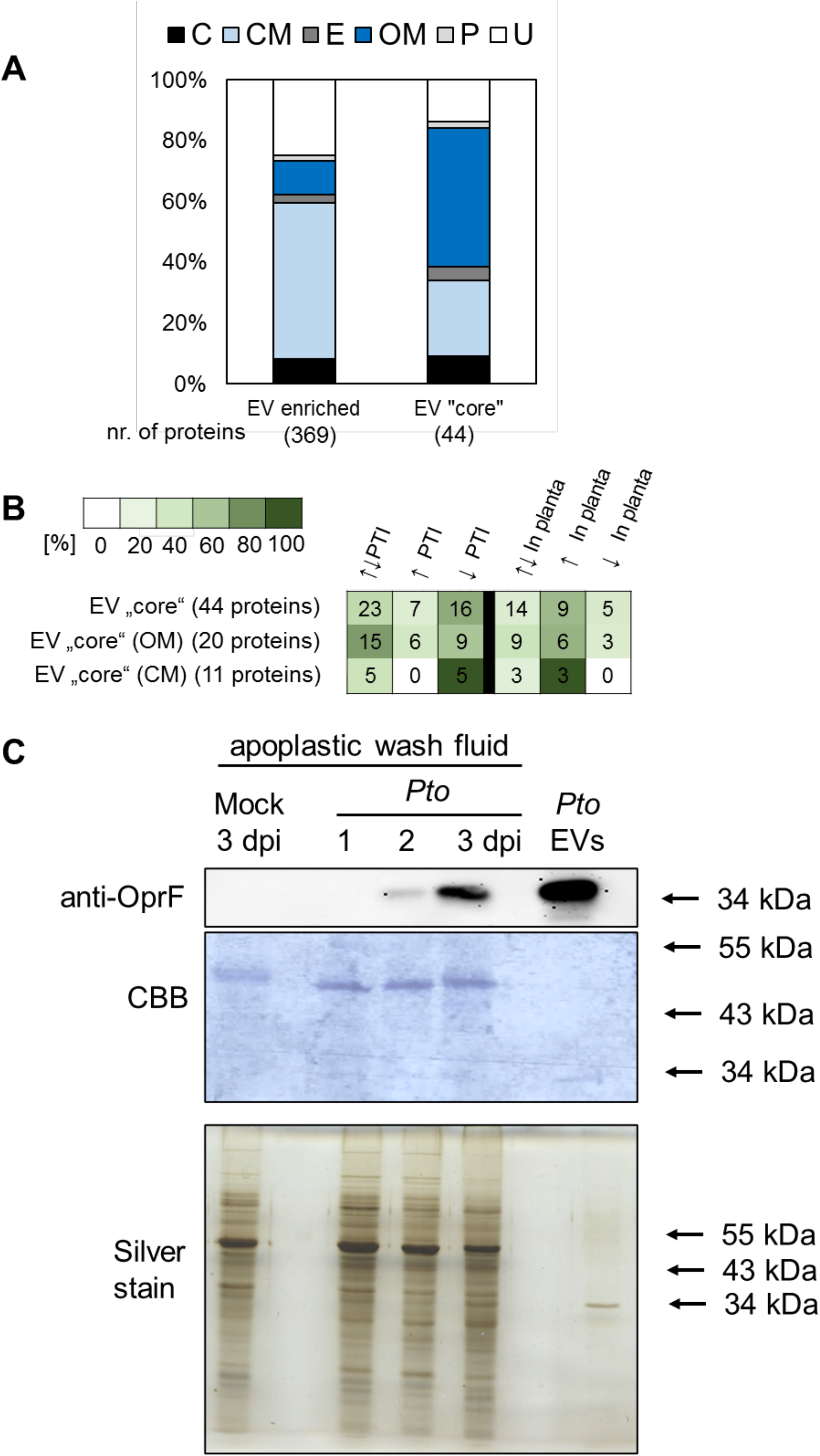
Prediction of Pseudomonas “core” EVs proteins identifies biomarkers for the detection of *Pto* DC3000 EVs *in planta*. **A)** Predicted protein localization of the *Pto* DC3000 EV enriched proteins whose orthologs are found in *P. aeruginosa* EV proteomes [%] total number of identified proteins is 44. **B)** Heat-map representing transcriptional changes in genes encoding *Pseudomonas* “core” EV proteins, EV “core” proteins localized on outer membrane (OM) and EV “core” proteins localized on cytoplasmic membrane (CM). The intensity of green colour represents percentage of affected genes and the numbers in boxes show the exact number of affected genes. **C)** Immunoblot monitoring OprF, a predicted *Pto* DC3000 EV biomarker, in EVs collected from apoplastic fluids. OprF antibodies detect bands in apoplastic fluids from *A. thaliana* infected with *Pto* DC3000 and in gradient enriched EVs. Coomassie Brilliant Blue (CBB) and silver staining are shown as loading control.

**Table 2.**
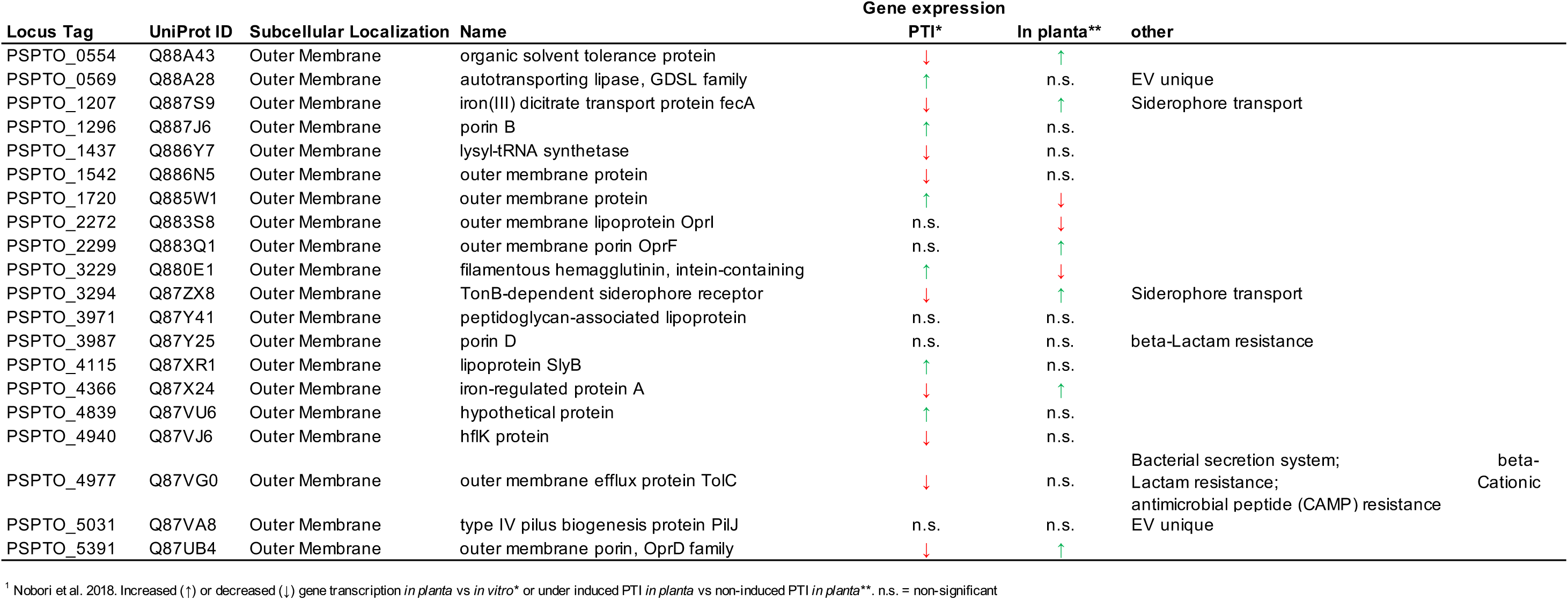
*In planta* transcription^1^ of genes coding proteins suggested as promising candidates to be EV markers

One of the predicted EV markers is OprF (Outer membrane porin OprF; Fig. 5C black labelling), which we used for immunodection of *Pto* DC3000 EVs *in planta* and purified from *Pto* DC3000 cultures. Using *anti*-OprF antibodies, we identified specific bands in filtered apoplastic fluids of *A. thaliana* leaves infected with *Pto* DC3000 at two and three days post infection but not in control treated plants (Fig. 8C). This provides additional evidence that *Pto* DC3000 releases EVs *in planta* during infection.

## Discussion

Bacterial EVs have been widely studied for their content and biological importance in various human diseases. Until recently, however, EV production by phytopathogenic bacteria has been mostly disregarded (Rybak & Robatzek, 2019). In the past few years, the attention in plant-microbe interactions has turned slowly towards EV signalling. EV production by phytopathogenic bacteria was shown for *A. tumefaciens* C58, *Xanthomonas campestris* pv *campestris* (*Xcc*), *X. campestris* pv *vesicatoria, Pto* T1, *Pto* DC3000 and *Xyllela fastidiosa* subsp. *fastidiosa* Temecula-1, subsp. *pauca* 9a5c and subsp. Fb7 (Bahar *et al.*, 2016, Feitosa-Junior *et al.*, 2019, Chowdhury & Jagannadham, 2013, Ionescu *et al.*, 2014, Knoke *et al.*, 2020, McMillan *et al.*, 2020, Sidhu *et al.*, 2008, Sole *et al.*, 2015). In this study, we provide several pieces of evidence for the role of EVs in the interaction of *Pto* DC3000 with plants: i) the bacteria produce EVs *in planta* during infection; ii) proteins responding to the plant environment are enriched in EVs; iii) known MAMPs and effectors are associated with EVs; and iv) plants respond to EVs with prototypic PTI reactions.

We show that *Pto* DC3000 produces spheres bulging from its outer membrane and releases spherical vesicles into the environment (Fig. 1A, 7B, 7C). Our data indicate the production of EVs predominantly in the form of OMVs albeit the predicted localization profiles of the EV-enriched proteins also suggest OIMV production (Fig. 7A, 7B, 7C, 8A), both representing well-established routes of vesicle release in Gram-negative bacteria (Toyofuku *et al.*, 2019). The more polydisperse size seen for EVs recovered from apoplastic fluids of susceptible *Pto* DC3000 infected plants could not only represent a mixed pool of plant and bacterial derived EVs, but also suggests that *Pto* DC3000 might release EVs from different biogenesis routes including membrane blebbing and explosive cell lysis.

Our results suggest the regulated production of EVs and the specific enrichment of proteins to EVs. *Pto* DC3000 cultures produced more EVs with increasing bacterial density during exponential-phase growth, showing a correlation between bacteria and EV numbers (Fig. 1E). The production of EVs by *Pto* DC3000 was slightly responsive to bacterial growth style and isolation technique. EVs produced from biofilm *Pto* DC3000 and fluid planktonic culture were smaller and more negatively charged than gradient enriched planktonic EVs. Moreover, heat inactivation of bacteria increased vesicle numbers and size (Fig. 1D, 1F). Since EVs from heat inactivated bacteria did not differ in charge profiles compared to untreated EVs (Fig. S6D), it is possible that explosive cell lysis contributes to the higher EV numbers (Toyofuku *et al.*, 2019). Turnbull *et al.* demonstrated that explosive cell lysis of a sub-population of cells from *P. aeruginosa* biofilms results in the generation of bacterial EVs (Turnbull *et al.*, 2016). Furthermore, heat shock stimulates the release of OMVs, most likely a result of high levels of un- and misfolded proteins accumulating in heat-stressed cells (Macdonald & Kuehn, 2013, McBroom & Kuehn, 2007). In this context it is interesting to note that the biophysical characteristics of *Pto* DC3000 EVs remained largely unchanged under disruptive treatment conditions except boiling (Fig. 2). Thus, the EVs from *Pto* DC3000 are stable and can maintain functionality as reported previously for other bacteria (Alves *et al.*, 2016, Arigita *et al.*, 2004, Frank *et al.*, 2018).

A range of activities has been associated with bacterial EVs. This includes modulation of host immunity i.e. through EVs presenting MAMPs and delivering effector molecules (Bahar *et al.*, 2016, Schwechheimer & Kuehn, 2015). We found that EVs from *Pto* DC3000 cultures elicited a robust induction of the *FRK1* defence marker gene (Fig. 4B, 4C, 4E). Compared with EVs from *Acidovorax* and *Xanthomonas* bacteria, *Pto* DC3000 EVs provoked a modest induction of defence gene expression (Bahar *et al.*, 2016). Despite this, the *Pto* DC3000 EVs did not trigger a growth arrest in Arabidopsis seedlings (Fig. 4D). These results suggest that the immunogenic activities of the EVs, but not a longer-term trade-off with growth, may be relevant in their interaction with plants. In contrast to our results, McMillan et al. reported significant seedling growth repression in response to *Pto* DC3000 EVs (McMillan *et al.*, 2020). This disparity in results may be due to several factors, including differences in the growth conditions of both the bacterial cultures and the *A. thaliana* seedlings, the type of biochemical isolation of EVs and vesicle dose. That EVs can serve as protective vaccines has been reported for many bacteria but not phytopathogens. We show here that plants pre-treated with *Pto* DC3000 EVs were modest, yet significantly protected against subsequent infection with *Pto* DC3000 bacteria (Fig. 4A). Consistently, EV pre-treatments did not inhibit a MAMP-induced ROS burst (Fig. 6A). The stronger protective immune response observed by McMillan et al. may be due to differences in experimental procedures, as noted above (McMillan *et al.*, 2020). In addition, McMillan et al. showed that pre-treatment with bacterial EVs provided protection against subsequent oomycete infection (McMillan *et al.*, 2020). The potential for *Pto* DC3000 EVs to induce broad-spectrum resistance supports a role for PTI.

PTI responses to *Pto* DC3000 involve recognition by FLS2, EFR and LIPO-OLIGOSACCHARIDE-SPECIFIC REDUCED ELICITATION (LORE), which detect immunogenic flg22, elf18 and 3-OH-FAs (Wan *et al.*, 2019). EVs from bacterial phytopathogens are enriched in EF-Tu and LPS (Bahar *et al.*, 2016, Feitosa-Junior *et al.*, 2019, Sidhu *et al.*, 2008), suggesting the presence of elf18 and 3-OH-FAs. Bahar *et al.* demonstrated that BRI1-ASSOCIATED KINASE 1 (BAK1) and SUPPRESSOR OF BIR 1 (SOBIR1), interacting co-receptors of PRRs, mediate the immunogenic perception of EVs from *X. campestris* pv. *campestris* (Bahar et al., 2016). We show that our vesicle samples from *Pto* DC3000 elicit immune responses that are dependent on FLS2 (Fig. 4E). Despite depleting extracellular components from EV samples by density gradient centrifugation, it is possible that flagella co-purify with the *Pto* DC3000 EV samples, since filamentous structures were observed in SEM analysis (Fig. 1B). Contamination of flagella in EVs was reported to contribute to the detection of FliC in EVs from *P. aeruginosa* (Bauman & Kuehn, 2006). However, flagella proteins such as FliC have a specific affinity for EVs and are involved in EV production in *Escherichia coli* (Manabe et al., 2013). We cannot exclude that flagella proteins play roles in EV production in *Pto* DC3000, evidenced by the finding that six flagella-associated proteins were enriched in *Pto* DC3000 EVs (Table S1).

Why *Pto* DC3000 produces EVs during infection is unclear. It is evident that the plant apoplast represents a stressful environment for its colonizing bacteria. Bacteria respond to environmental stress with the production of EVs, which allows for cell surface remodelling, secretion of degraded and damaged cargo, and uptake of nutrients in bacterial communities i.e. by packaging transporters in EVs (Schwechheimer & Kuehn, 2015, Toyofuku *et al.*, 2019, Zingl *et al.*, 2020). As the growth state of bacteria determines the nutrient availability, the production of EVs could support the growth of *Pto* DC3000 in culture and *in planta.* Proteomics analysis of gradient enriched *Pto* DC3000 EVs identified 890 vesicle-associated proteins, of which 369 were enriched in the EVs relative to the cellular proteome (Fig. 5, Table S1). The mechanisms by which this enrichment occurs suggests a selective delivery of cargo into the EVs and should be investigated in the future. Of the EV-enriched proteins, six out of ten GO biological process categories were related to transport mechanisms (Fig. 6B). Proteins involved in siderophore transport were enriched in the EVs and have recently been shown to play a role in *Pto* DC3000 infection success (Nobori *et al.*, 2018). Interestingly, the expression of genes coding for all siderophore transport proteins enriched in EVs was upregulated *in planta* compared to *in vitro* conditions as well as downregulated upon induction of PTI (Fig. 6B; Table 1) (Nobori *et al.*, 2018). Thus, regulation of siderophore transport proteins can be considered as an adaptive response of *Pto* DC3000 to iron/metal ion availability, and secretion into EVs may allow improved acquisition of iron, analogous to EV secretion of the siderophore mycobactin in *Mycobacterium tuberculosis* (Prados-Rosales et al., 2014). The plant’s apoplast, which is the niche colonized by *Pto* DC3000 represents an environment where bacteria are challenged with iron acquisition and plant defence molecules (Nobori *et al.*, 2018). The role of bacterial EVs in metal acquisition is not restricted to iron. *Neisseria meningitidis* produces OMVs, which are enriched in zinc acquisition proteins (Lappann *et al.*, 2013), and zinc regulates siderophore biosynthesis genes in *Pseudomonas fluorescens* (Rossbach et al., 2000). It is thus feasible that *Pto* DC3000 produces EVs to help it adapt to metal conditions in the environment including the plant’s apoplast.

The hypothesis that *Pto* DC3000 uses EVs to adapt to the growth environment is supported by our finding that *Pto* DC3000 EVs are enriched in proteins related to the KEGG categories ß-lactam resistance and cationic antimicrobial peptide resistance (Fig. 6B). Several studies demonstrated that EVs can improve bacterial survival during antibiotic exposure. *Stenotrophomonas maltophilia* produced more EVs upon treatment with the ß-lactam antibiotic imipenem (Devos *et al.*, 2015, Devos *et al.*, 2017). Its EVs contained ß-lactamase and increased *S. maltophilia* survival in the presence of antibiotics (Devos *et al.*, 2017). Plants defend infection by upregulation of many defence-related gene including genes coding for antimicrobial peptides (Campos *et al.*, 2018). It is possible that *Pto* DC3000 produces EVs to counter the action of plant-derived antimicrobial peptides. Collectively, we propose that *Pto* DC3000 produces EVs to improve its growth capacity both in culture and *in planta*. These findings should stimulate further studies on the role of EVs in the interaction of bacteria with plants, for example identifying the composition of EVs *in planta* using biomarkers.

## Experimental procedures

### Bacterial strains and growth

*Pseudomonas syringae* pv. *tomato* DC3000 (*Pto* DC3000) used in this study were routinely cultured at 28 °C in King’s B medium containing 50 μg/mL Rifampicin at 180 rpm and on plates with 1% agar without agitation. Planktonic growth was performed in 500 mL and growth rates were measured over time as OD_600_. Biofilm growth was measured after 24 h on plate, transferring all bacteria per plate in 10 mL 0.85% saline.

### Plant material and growth conditions

*Arabidopsis thaliana* ecotype Columbia (Col-0),*pFRK1::GUS* (Kunze et al. 2004),*fls2c* (Zipfel et al. 2004) and *efr-1* (Zipfel et al. 2006) mutants were used in this study. For bacterial infections and ROS assays, Col-0 plants were soil-grown at 21–22 °C and 8 h photoperiod. For GUS assays, RT-qPCR analysis and induced growth arrest, seedlings were sterile grown on Murashige and Skoog (MS) plates supplemented with 1% sucrose and 1.5% gelrite (Duchefa, Netherlands) pH 5.8 for four days (after 2-4 days stratification in the dark at 4 °C), then transferred to 96-well plates containing 150 μL ½ MS medium supplemented with 1% sucrose per well and grown for eleven to twelve days in at 22 °C and 16 h photoperiod (120 – 150 μE.m^-2^.s^-1^).

### Extraction and purification of bacterial EVs

EVs were routinely isolated from planktonic cultures at early-logarithmic to late-stationary phases as well as biofilm cultures (Fig. S2B, S2D). 100 mL of planktonic grown bacteria and 10 mL of biofilm grown bacteria, respectively, were pelleted at 4,500 x g for 2 x 20 min, the supernatant was decanted and passed through a 0.22 μm membrane (fluid samples; Fig. S2A). Particles were pelleted from the cell-free supernatant at 100,000 x g for 1.5 h. The pellet was resuspended in 1.7 mL 1mM EDTA and loaded on sucrose density step-gradient (1.7 mL of sucrose 25%, 35%, 45%, 50%, 55%) and centrifuged at 160,000 x g for 18 h. 2 mL samples were collected from each of the sucrose density steps and diluted with 1 mM EDTA to 30 mL. Particles were pelleted at 100,000 x g for 2 h and the pellets were each resuspended in 0.16 mL 1 mM EDTA (gradient enriched samples; Fig. S2A). EV samples were immediately frozen in liquid nitrogen. Since most EVs migrated to the 55% density fraction (Fig. S6A), we then collected EVs across fractions 3 to 5, which were less variable in ζ-potential and size compared to fractions 1 and 2 (Fig. S6B, S6C).

### Extraction of leaf apoplastic fluids

Apoplastic fluids were collected from leaves of six to seven weeks old plants. The leaves were cut of the rosette and vacuum infiltrated with particle-free 1 mM EDTA. After removing excess buffer, infiltrated leaves were placed into 20 mL syringes and centrifuged in 50 mL conical tubes at 900 x g for 20 minutes at 4°C. The resulting apoplastic wash was passed through a 0.22 μm membrane (apoplastic fluid samples).

### EV quantification, size and charge measurements

EVs were quantified, size and charge measured by Nanoparticle Tracking Analysis (NTA) using ZetaView^®^ BASIC PMX-120 (Particle Metrix, Germany) at room temperature. To detect EVs, we used the manufacturer’s default settings for liposomes. Particle quantification and size measurements were performed by scanning eleven cell positions each and capturing 30 frames per position with the following settings: Focus: autofocus; Camera sensitivity for all samples: 85; Shutter: 100; Scattering Intensity: detected automatically. After capture, the videos were analysed by the in-built ZetaView Software 8.05.11 [ZNTA] with the following specific analysis parameters: Maximum area: 1000, Minimum area 5, Minimum brightness: 25, Tracelength: 15 ms. Hardware: embedded laser: 40 mW at 488 nm; camera: CMOS. For particle charge measurements, the same settings were used except Minimum brightness: 30. Statistical analysis was performed using either One-way ANOVA with Tukey post hoc test or Welsch’s ANOVA with Dunnett’s T3 multiple comparisons post hoc test.

All samples were diluted in particle-free 1 mM EDTA buffer, checked with NTA. Unconditioned King’s B medium contained up to 1.4x 10^9^ particles (Fig. 1D). *Pto* DC3000 cultures contained increasing particles numbers with cultivation time: ≈ 2.8x 10^9^ particles at OD_600_ = 3-4 (50% of influence); ≈ 3.7x 10^9^ particles at OD_600_ = 4.5-5.5 (37% of influence), ≈ 7*10^9^ particles at OD_600_ 7.5-9 (20% of influence), and ≈1.1x 10^10^ particles at OD_600_ 10 −11 (13% of influence) (Fig. 1D). We therefore focused our measurements on samples collected from OD_600_ > 7.5, which shows lower than 20% influence of particles from the medium (Fig. 1D), as well as calculated EV concentrations to the colony forming units (CFU) of the bacterial cultures.

### Scanning electron microscopy

Planktonic grown bacteria at OD_600_ = 3-4 and gradient enriched EVs (0.5 to 1.5x 10^10^ particles) were used for scanning electron microscopy (SEM). The cells were chemically fixed using 2.5% glutaraldehyde in 50 mM cacodylate buffer (pH 7.0) containing 2 mM MgCl2. Then the cells were applied to a glass slide, covered with a cover slip and plunge frozen in liquid nitrogen. After this, the cover slip was removed and the cells were place in fixation buffer again. After washing 4 times with buffer, post-fixation was carried out with 1% OsO4 for 15 min. Two additional washing steps with buffer were followed by three times washing with double distilled water. The samples were dehydrated in a graded acetone series, critical-point-dried and mounted on an aluminium stub. To enhance conductivity, the samples were sputter-coated with platinum. Microscopy was carried out using a Zeiss Auriga Crossbeam workstation at 2 kV (Zeiss, Oberkochen, Germany). The vesicle size was manually measured across five randomly selected SEM micrographs using Fiji software (Schindelin *et al.*, 2012).

### Transmission electron microscopy

Planktonic grown *Pto* DC3000 at OD_600_ = 3-4 were used for ultrathin sectioning and subsequent transmission electron microscopy (TEM). The cells were concentrated by centrifugation and the cells were high-pressure frozen using a Leica HPM100 (Leica Microsystems, Wetzlar, Germany). This was followed by freeze-substitution with 0.2% osmium tetroxide, 0.1% uranyl acetate, 9.3% water in water-free acetone in a Leica AFS 2 (Leica Microsystems, Wetzlar, Germany) as described previously (Flechsler *et al.*, 2020). After embedding in Epon 812 substitute resin (Fluka Chemie AG, Buchs Switzerland), the cells were ultrathin sectioned (50 to 100 nm thickness) and post-stained for 1 min with lead citrate. Transmission electron microscopy of ultrathin sections was carried out with a JEOL F200 cryo-S(TEM), which was operated at 200 kV and at room temperature in the TEM mode. Images were acquired using a bottom-mounted XAROSA 20 mega pixel CMOS camera (EMSIS, Münster, Germany).

### *Pto* DC3000 infection assay

Overnight plate-grown *Pto* DC3000 cells were resuspended in 10 mM MgCl2 and diluted to OD_600_ = 0.0006. Using a needle-less syringe, the bacterial suspension was infiltrated into mature leaves of five to six weeks old plants, three leaves per plant. For pre-treatments, gradient enriched EVs from planktonic *Pto* DC3000 (concentration ≈ 1.10^10^), and 0.02 mM EDTA as a negative control and 100 nM flg22 (EZbiolabs) as a positive control were syringe-infiltrated into leaves 24 h prior *Pto* DC3000 inoculation. Discs of the infected leaves (one disc per leaf, 0.6 cm diameter) were excised at one, two- or three-days post infection (dpi). The three leaf discs from each plant were pooled and ground in 1 mL 10 mM MgCl2. Serial dilutions were plated on LB medium with rifampicin (50 μg/mL) and bacterial colonies were counted one day after incubation at 28 °C. Statistical analysis was performed using two tailed Welsch’s t-test.

### Histochemical β-glucuronidase (GUS) staining

The histochemical GUS assay was performed with eleven day old seedlings. Seedlings were treated with gradient enriched *Pto* DC3000 EVs (concentration ≈ 1.10^10^), 100 nM flg22 (EZbiolabs) or as a control with 0.02 mM EDTA for 18 h. Treated seedlings were immersed in X-Gluc buffer [2 mM X-Gluc (Biosynth), 50 mM NaPO4, pH 7, 0.5 % (v/v) Triton-X100, 0.5 mM K-ferricyanide] for 16 h at 37 °C. Chlorophyll was removed by repeated washing in 80 % (v/v) ethanol. Observations were made on a WHX 6000 digital microscopy (Krckova *et al.*, 2018).

### Fluorimetric GUS assay

For fluorimetric GUS assays, eleven to twelve days old seedlings were treated with gradient enriched *Pto* DC3000 EVs (concentration ≈ 1.10^10^) or with 100 nM flg22 (EZbiolabs) or as a control with 0.02 mM EDTA for 18 h. Treated seedlings were frozen in liquid nitrogen in 2 mL conical tubes containing 2 clean sterile glass beads and liquid nitrogen. The frozen samples were dry homogenized using a Retch mixer mill (Retch). Homogenized samples were kept on ice and cold (4 °C). For total protein extraction, GUS extraction buffer was added as described (Andriankaja *et al.*, 2007) [50 mM sodium phosphate (pH 7); 10 mM 2-mercaptoethanol; 10 mM Na2EDTA; 0.1% Triton X-100; 0.1% sodium lauryl-sarcosine and PPIC]. GUS activities were measured fluorimetrically in reaction buffer (see below) using Methylumbelliferyl-β-D-glucuronic acid dihydrate (MUG), (Biosynth) as a substrate. Reaction buffer was the same solution as extraction buffer with one modification: PPIC was replaced by 1 mM MUG. The fluorescence was measured using TECAN fluorimeter at excitation 360 nm and emission 465 nm. The enzymatic activity of the sample was calculated to protein concentration measured by Bradford protein assay. The absorbance was measured using TECAN spectrometer absorbance at 595 nm. Statistical analysis was performed using One-way ANOVA with Tukey post hoc test.

### RNA extraction and RT-qPCR analysis

Gene transcription analysis was performed with twelve days old seedlings. The seedlings were treated with gradient enriched EVs (concentration 1.10^10^) and 0.02 mM EDTA as control for 3 h, frozen in liquid nitrogen and ground with 2.5-mm diameter silica beads using a homogenizer (Retch, Germany). Total RNA was isolated using a TRIzol^®^ reagent (Invitrogen, USA) according to the manufacturer’s protocol. The extracted RNA was treated with a DNA-free kit (Ambion, USA). Subsequently, 1 μg of RNA was converted into cDNA with M-MLV RNase H–Point Mutant reverse transcriptase (Promega Corp., USA) and an anchored oligo dT21 primer (Metabion, Germany). Gene transcription was quantified by qPCR using a LightCycler 480 SYBR Green I Master kit and LightCycler 480 (Roche, Switzerland). The PCR conditions were 95 °C for 10 min followed by 45 cycles of 95 °C for 10 s, 55 °C for 20 s, and 72 °C for 20 s. Melting curve analyses were then carried out. Relative transcription was normalized to the housekeeping gene *AtTIP41* (Czechowski *et al.*, 2005). Primers were designed using PerlPrimer v1.1.21 (Marshall, 2004). The primers used are *AtFRK1*_FP, GCCAACGGAGACATTAGAG; *AtFRK1 _RP*, CCATAACGACCTGACTCATC. Statistical analysis was performed using One-way ANOVA with Tukey post hoc test.

### Seedling growth analysis

Four days old seedlings were transferred from MS solid media into the liquid MS media in transparent 96-well microplates. Each well contained 100 μL of media either containing 0.02 mM EDTA as a control or gradient enriched *Pto* DC3000 EVs (concentration ≈ 1.10^10^) or with 100 nM flg22 (EZbiolabs) as a positive control. After eight days, the treated seedlings were dried using paper towel and then the fresh weight was measured. Based on the weight of each seedling relative seedlings growth [%] to control seedlings was calculated. Statistical analysis was performed using Welsch’s ANOVA with Dunnett’s T3 multiple comparisons post hoc test two tailed Student t-test.

### ROS measurements

ROS production was determined using the luminol-based assay as previously described (Mersmann *et al.*, 2010). Briefly, leaves of five to six weeks old *A. thaliana* plants were infiltrated with gradient enriched EVs (concentration ≈ 1.10^10^). After 2 h, discs were excised from the infiltrated leaves and 24 h incubated in ddH2O at 22 °C. Then, the leaf discs were treated with 100 nM flg22 or 100 nM elf18 (EZbiolabs) to induce the production of ROS. The total photon count was collected for 45 min using a TECAN luminometer. Statistical analysis was performed using two tailed Student t-test.

### Proteomics

We isolated proteins from *Pto* DC3000 whole cell lysates (WC) (Park *et al.*, 2014) and outer membrane (OM) (Choi *et al.*, 2011) as previously described. Briefly, WC and OM isolated from *Pto* DC3000 liquid culture (OD_600_ = 3-4). The cells were pelleted via centrifugation (12,000 x g for 10 min). For WC the pellet was resuspended in 1 mL of 20 mM Tris-HCl (pH 8.0), frozen in liquid nitrogen, three times thawing-freezing, and three times sonicated for 10 min at 4 °C. The samples were centrifuged at 6,000 x g for 10 min at 4 °C and supernatants were collected and frozen in liquid nitrogen. For OM preparations, the pellet was resuspended in 1 mL 20 mM Tris-HCl (pH 8.0), sucrose (20%), followed by adding 5 uL Lysozyme (15 mg/mL) and 10 μl 0.5 M EDTA, incubation for 40 min on ice and adding 20 μL 0.5 M MgCl2. After centrifugation at 9,500 x g for 20 min at 4°C, the pellet was resuspended in 1 mL ice-cold 10 mM Tris-HCl (pH 8.0) followed by sonication three times for 10 min on ice. The samples were then centrifuged at 8,000 x g for 5 min at 4 °C, washed with cold 10 mM Tris-HCl (pH 8.0), resuspend in cold, sterile MilliQ water followed by three times freezing-thawing in liquid nitrogen, incubation for 20 min at 25 °C and adding the sarcosyl to final concentration 0.5%. The samples were then centrifuged at 40,000 x g for 90 min at 4 °C, the pellet was resuspended in ice-cold 10 mM Tris-HCl (pH 8.0) and frozen in liquid nitrogen. Gradient enriched EVs were isolated as above described (Fig. S1C). For proteomics, the samples were denatured by addition of 1 x SDS loading buffer. In-gel trypsin digestion was performed according to standard procedures (Shevchenko *et al.*, 2006). Briefly, 2 μg of EV and OM samples and 20 μg of WC samples were loaded on a NuPAGE 4-12% Bis-Tris Protein gels (Thermofisher Scientific, US), and the gels were run for 3 min only. Subsequently, the still not size-separated single protein band per sample was cut, reduced (50 mM DTT), alkylated (55 mm CAA, chloroacetamid) and digested overnight with trypsin (trypsin-gold, Promega).

### LC-MS/MS data acquisition

Peptides generated by in-gel trypsin digestion were dried in a vacuum concentrator and dissolved in 0.1% formic acid (FA). LC-MS/MS measurements were performed on a Fusion Lumos Tribrid mass spectrometer (Thermo Fisher Scientific) equipped with an Ultimate 3000 RSLCnano system. Peptides were delivered to a trap column (ReproSil-pur C18-AQ, 5 μm, Dr Maisch, 20 mm × 75 μm, self-packed) at a flow rate of 5 μL/min in 100% solvent A (0.1% formic acid in HPLC grade water). After 10 min of loading, peptides were transferred to an analytical column (ReproSil Gold C18-AQ, 3 μm, Dr Maisch, 400 mm × 75 μm, self-packed) and separated using a 50 min gradient from 4% to 32% of solvent B (0.1% formic acid in acetonitrile and 5% (v/v) DMSO) at 300 nL/min flow rate. Both nanoLC solvents contained 5% (v/v) DMSO.

The Fusion Lumos Tribrid mass spectrometer was operated in data dependent acquisition and positive ionization mode. MS1 spectra (360–1300 m/z) were recorded at a resolution of 60,000 using an automatic gain control (AGC) target value of 4e5 and maximum injection time (maxIT) of 50 ms. After peptide fragmentation using higher energy collision induced dissociation (HCD), MS2 spectra of up to 20 precursor peptides were acquired at a resolution of 15.000 with an automatic gain control (AGC) target value of 5e4 and maximum injection time (maxIT) of 22 ms. The precursor isolation window width was set to 1.3 m/z and normalized collision energy to 30%. Dynamic exclusion was enabled with 20 s exclusion time (mass tolerance +/-10 ppm).

### Computational analysis of proteomes

LFQ values were used in the statistical analysis of proteome data. To select EV-enriched proteins, Welch t-test were used to compare protein intensities between EV and WC samples. The resulted p-values were corrected using the Benjamini-Hochberg method to control the false discovery rate (FDR). The proteins with FDR < 0.05 and with the intensity in EV at least twice higher than in WC were selected as EV-enriched proteins (n = 207). In addition, we selected proteins that were exclusively identified in at least three (out of four) replicates of EV. (n = 162). A complete list of EV-enriched proteins is given in Table S1. The functional enrichment analysis of the EV proteins were performed using the DAVID functional annotation tool (Huang da *et al.*, 2009a, Huang da *et al.*, 2009b).

### Database searches

Peptide identification and quantification was performed using MaxQuant (version 1.6.3.4) with its built-in search engine Andromeda (Cox *et al.*, 2011, Tyanova *et al.*, 2016). MS2 spectra were searched against a *Pseudomonas syringae pv tomato* protein database (UP000002515, downloaded from Uniprot 04.05.2020) supplemented with common contaminants (built-in option in MaxQuant). For all MaxQuant searches default parameters were employed. Those included carbamidomethylation of cysteine as fixed modification and oxidation of methionine and N-terminal protein acetylation as variable modifications. Trypsin/P was specified as proteolytic enzyme. Precursor tolerance was set to 4.5 ppm, and fragment ion tolerance to 20 ppm. Results were adjusted to 1% false discovery rate (FDR) on peptide spectrum match (PSM) and protein level, employing a target-decoy approach using reversed protein sequences. Label-free quantification (LFQ algorithm) was enabled. The minimal peptide length was defined as 7 amino acids and the “match-between-run” function was not enabled. Each sample type (EV, OM, WC) was analysed in biological quadruplicates (Table S1).

We used available localization prediction data at pseudomonas genome database (pseudomonas.com) (Winsor *et al.*, 2016). Predicted protein localizations are presented as stacked bar charts (made in MS Excel) as percentage to total number of the proteins in analyzed sample. We used the available software DAVID bioinformatic resource 6.8 (https://david.ncifcrf.gov/) for GO term and KEGG pathway analysis, and the adjusted p-value cut-off was set to 0.05 (Huang da *et al.*, 2009a, Huang da *et al.*, 2009b). We compared the EV enriched proteins from *Pto* DC3000 with EV proteomes from planktonic grown *P. aeruginosa* PAO1 (Choi *et al.*, 2011, Park *et al.*, 2014, Reales-Calderon *et al.*, 2015). We focussed on the proteins that were identified in OMVs from *P. aeruginosa* PAO1 across all three studies and identified their gene orthologs in *Pto* DC3000 using the pseudomonas genome database (pseudomonas.com) (Winsor *et al.*, 2016). This set of proteins was compared to the *Pto*DC3000 EV-enriched proteins to predict EV biomarkers. The EV-enriched proteins were also compared with available *in planta Pto* DC3000 transcriptome and proteome data (Nobori *et al.*, 2018) (Nobori *et al.*, 2020).

### Immunoblot analysis

Standard immunoblot analysis was performed according to Sambrook at al. (1989). 10% SDS-PAGE gels were blotted onto PVDF Immobilon-P membranes (Millipore). *Pto* DC3000 OprF was detected using 1:2,000 diluted rabbit polyclonal antibody against OprF from *Pseudomonas aeruginosa* (Cusabio Biotech Co.). As secondary antibody, we used a 1:50,000 dilution of the anti-rabbit IgG-Peroxidase polyclonal antibody (Sigma-Aldrich, A0545). Signal detection was done using SuperSignal West FemtoMaximum Sensitivity Substrate (Pierce, Thermo Scientific), according to manufacturer’s instructions, and the images were captured using Vilber Lourmat Peqlab FUSION SL Gel Chemiluminescence Documentation System.

### Statistical analysis

Student *t*-test, Welsch’s *t*-test, One-way ANOVA followed by Tukey multiple comparisons test and Welsch’s ANOVA with Dunnett’s T3 multiple comparisons post hoc test were performed using GraphPad Prism version 8.3 for Windows, GraphPad Software, San Diego, California USA, www.graphpad.com

## Data availability

The mass spectrometry proteomics data have been deposited to the ProteomeXchange Consortium via the PRIDE (Perez-Riverol *et al.*, 2019) partner repository with the dataset identifier PXD023971.

## Author contributions

M.J. and S.R. designed research; M.J., C.L., K.R., C.M., L.B., B.S., A.B., A.K. performed research; M.J., C.L., K.R., C.M., A.B., A.K. and S.R. analysed data; E.S., J.S., F.M., J.M. developed protocols; M.J. and S.R. wrote the paper with inputs from all authors.

## Acknowledgements

We like to thank members of the Robatzek laboratory for fruitful discussions. We acknowledge Lucia Grenga and Catriona Thompson (JIC), Franziska Hackbarth and Hermine Kienberger (BayBioMS) for their laboratory assistance, Miriam Abele for her mass spectrometric support at the BayBioMS as well as Jennifer Grünert and Cornelia Niemann (LMU Biocenter) for technical assistance in electron microscopy. This research was supported by the Deutsche Forschungsgemeinschaft (to S.R.) through a Heisenberg fellowship (RO 3550/14-1) and the SFB924 “Yield” (TP B15). M.J. was supported by the European Structural and Investment Funds, OP RDE-funded project ‘CHEMFELLS4UCTP’ (no. CZ.02.2.69/0.0/0.0/17_050/0008485).

## Supplemental Information

**Figure S1. A)** The full-size SEM micrograph used in Fig. 1A of *Pto* DC3000 growth in planktonic culture (OD_600_ = 3-4). **B)** Size profile of EVs from *Pto* DC3000 planktonic cultures in fluid samples (OD_600_ = 7.5-11).

**Figure S2. Isolation of *Pto* DC3000 EVs. A)** Schematic overview of EVs isolation from planktonic cultures for fluid sample (1) and gradient enriched sample (2) analysis. **B)** Growth measurements of planktonic *Pto* DC3000 cultures. Orange indicates EV isolation from early exponential growth stages (OD_600_ = 1-2); green indicates EV isolation from late exponential growth stages (OD_600_ = 3-4). **C)** Schematic overview of EVs isolation from biofilm cultures. **D)** Growth measurements of biofilm *Pto* DC3000 cultures. The green dot represents the growth stage from which the bacteria were used for experiments.

**Figure S3. Biophysical parameters of particles in apoplastic fluids from *A. thaliana* plants infected with *Pto* DC3000. A, D, G)** Particle parameters over days post infection (dpi). **B, E, H)** Particle parameters in response to inoculation with different *Pto* DC3000 densities. **C, F, I)** Particle parameters in response to inoculation with different *Pto* DC3000 and co-treatment with flg22. Each dot represents value of independent samples for size and ζ-potential it represents median. 3-12 independent samples were used for each experiment. **J, K)** The profile of ζ-potential for each particle collected from apoplastic fluids of plants treated as indicated and gradient enriched EVs. Control = 0.2 mM EDTA; flg22 = 100 nM; n.t. = not treated; *Pto* DC3000 OD_600_ = 0.0006. Each treatment was 3 days long. The dots represent the mean across the ζ-potential values from independent samples: n = 8 (control); n = 10 (*Pto* DC3000); n = 6 (flg22); n = 4 (non-treatment).

**Figure S4. Pre-treatment with *Pto* DC3000 EVs induces resistance against subsequent *Pto* DC3000 infection. A)** Three individual biological repeats of *Pto* DC3000 growth (CFU) after infection into leaves of *A. thaliana* without and with EV pre-treatment at 3 dpi (control = 0.02 mM EDTA). Each biological repeat consists of 12 independent samples. **B)** *Pto* DC3000 growth (CFU) after infection (3 dpi) into leaves of *A. thaliana* without and with 100 nM flg22 1 day pre-treatment (mock = 10 mM MgCl2) n = 4. Asterisks represent the statistical difference between the treated and control samples (two tailed Student t-test p < 0.01).

**Figure S5. Characteristics of the proteomic analysis. A)** Barplot shows the number of identified proteins in each replicate. The solid line indicates the cumulative protein IDs and dashed line shows the shared protein IDs. **B)** Boxplot shows a comparable distribution of protein intensities from each replicate.

**Figure S6. Biophysical parameters of *Pto* DC3000 EVs across isolation methods. A-C)** NTA measurements of particle concentration (A), ζ-potential (B) and size (C) of *Pto* DC3000 EVs collected from each step of gradient enrichment. **D-E)** NTA analysis of particle ζ-potential (D) and size (E) of *Pto* DC3000 EVs from fluid samples before (live) and after boiling. Each dot represents an independent sample.

**Table S1.** Filtered proteomics data with proteins that were identified in at least in three out of four biological repeats in at least one variant (WC, OM and EV). Values used for volcano plot: Highlighted EV-enriched proteins. Subcellar localization of identified proteins. Flagellar proteins identified in EV enriched proteins.

**Table S2.** GO analysis of EV-enriched proteins

**Table S3.** Expression of EV-enriched proteins *in planta*

**Table S4.** *Pseudomonas* EV “core”

## Notes

### Competing Interest Statement

The authors have declared no competing interest.

