## Supplemental Figures S1-S6 for "Biophysical and proteomic analyses suggest functions of *Pseudomonas syringae* pv *tomato* DC3000 extracellular vesicles in bacterial growth during plant infection"

Figure S1

A

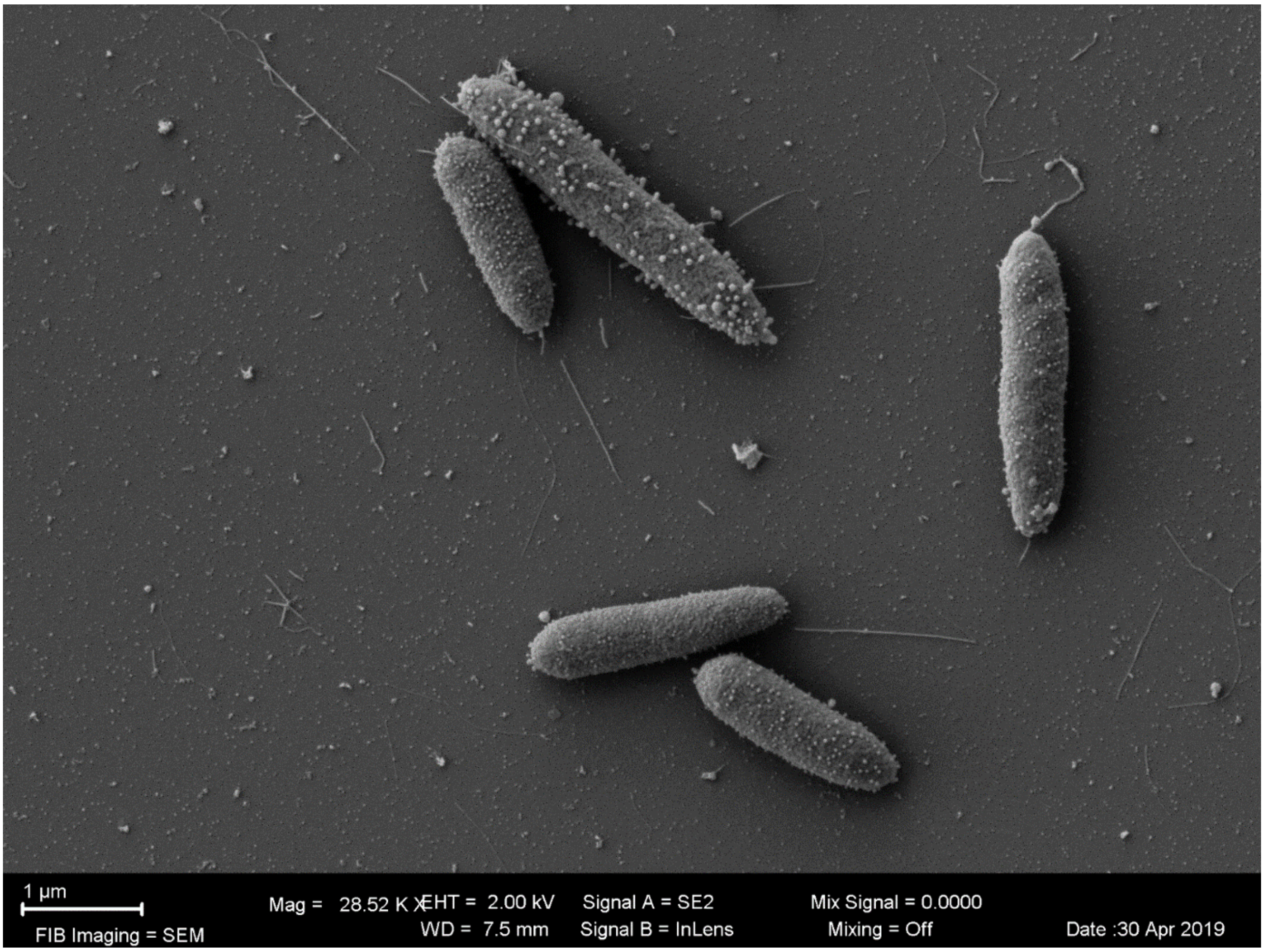


B

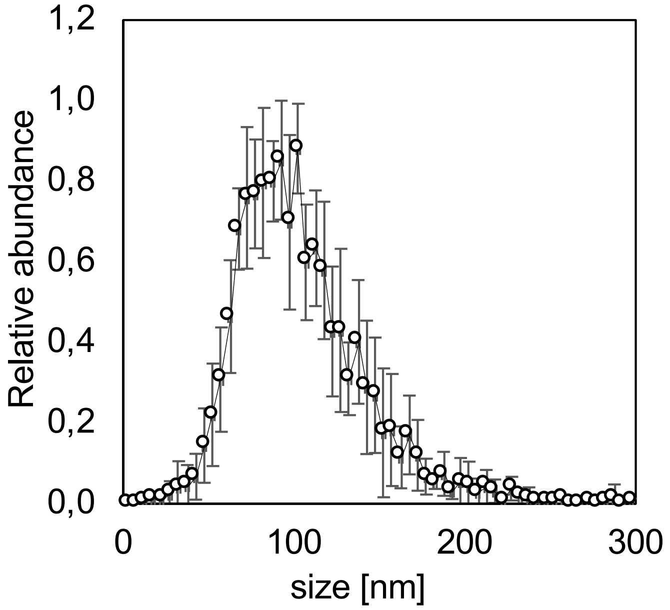


**Figure S1. A)** The full-size SEM micrograph used in Fig. 1A of *Pto* DC3000 growth in planktonic culture ($OD_{600}$ = 3-4). **B)** Size profile of EVs from *Pto* DC3000 planktonic cultures in fluid samples ($OD_{600}$ = 7.5-11).

Figure S2

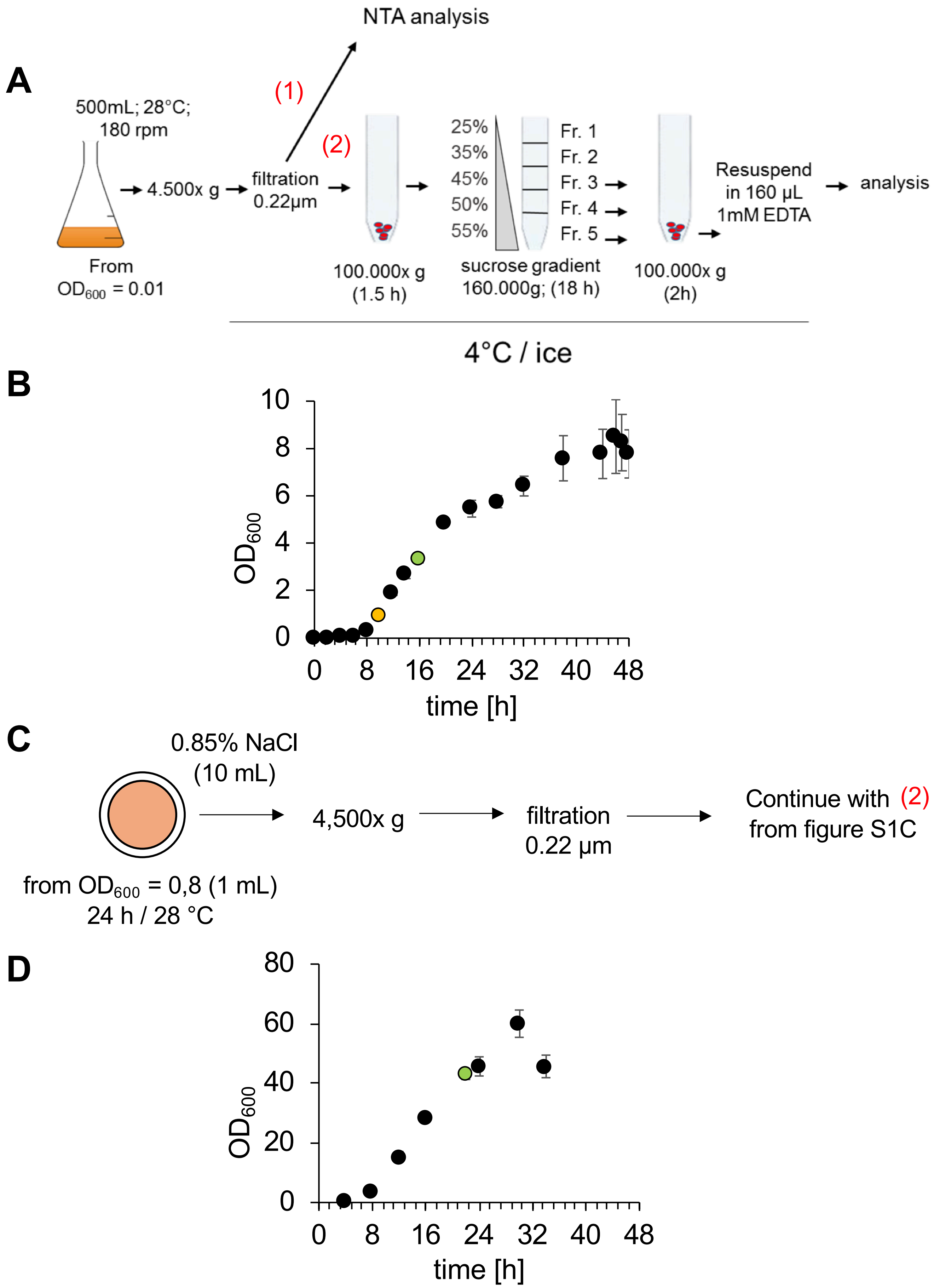


**Figure S2. Isolation of *Pto* DC3000 EVs. A)** Schematic overview of EVs isolation from planktonic cultures for fluid sample (1) and gradient enriched sample (2) analysis. **B)** Growth measurements of planktonic *Pto* DC3000 cultures. Orange indicates EV isolation from early exponential growth stages ($OD_{600}$ = 1-2); green indicates EV isolation from late exponential growth stages ($OD_{600}$ = 3-4). **C)** Schematic overview of EVs isolation from biofilm cultures. **D)** Growth measurements of biofilm *Pto* DC3000 cultures. The green dot represents the growth stage from which the bacteria were used for experiments.

Figure S3

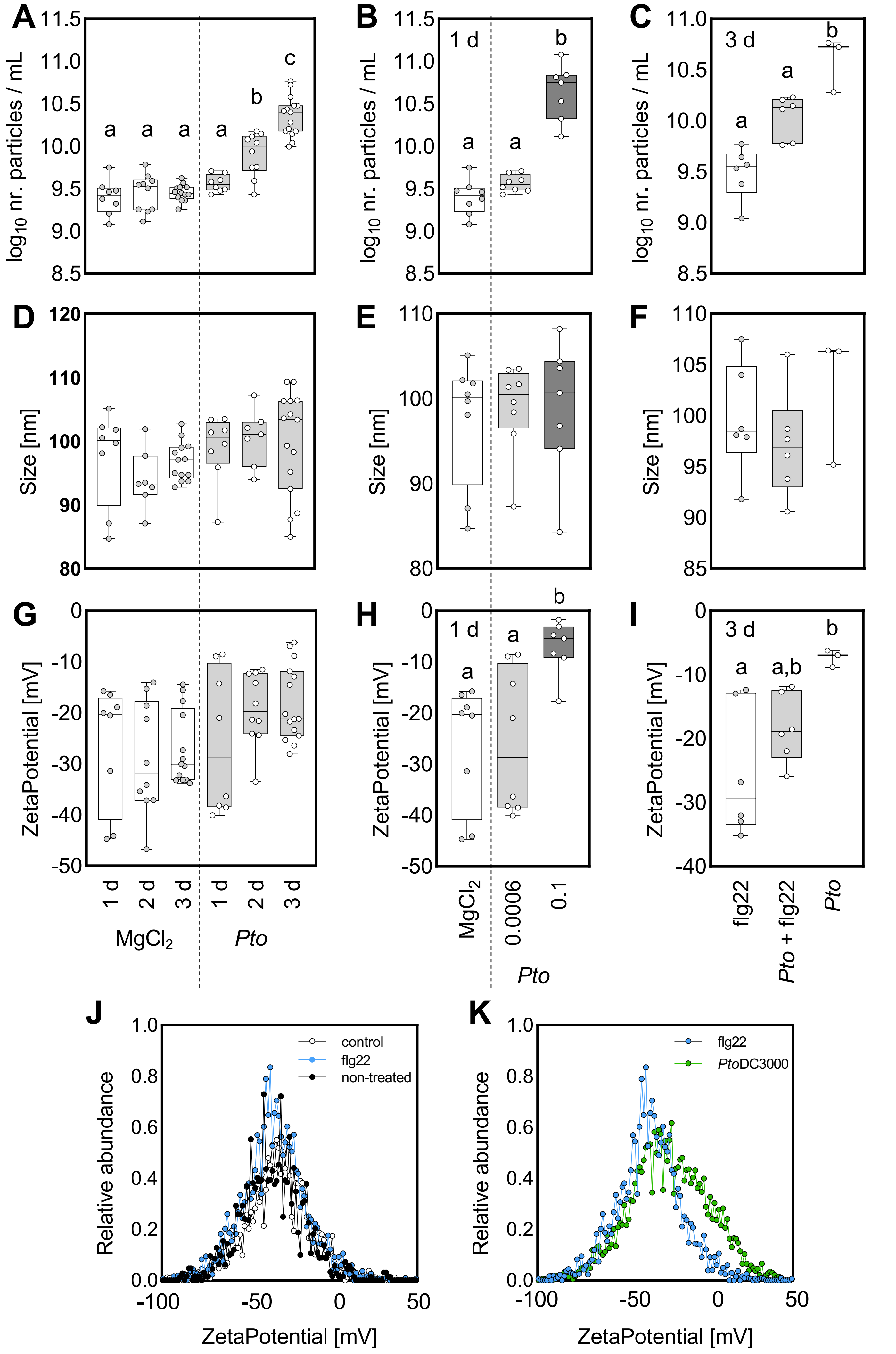

A
B
C
D
E
F
G
H
I
J
K
1 d
3 d
a
b
c
a,b
log10 nr. particles / mL
Size [nm]
ZetaPotential [mV]
1 d
2 d
3 d
MgCl2
Pto
0.0006
0.1
flg22
Pto + flg22
control
non-treated
PtoDC3000
Relative abundance

**Figure S3. Biophysical parameters of particles in apoplastic fluids from *A. thaliana* plants infected with *Pto* DC3000. A, D, G)** Particle parameters over days post infection (dpi). **B, E, H)** Particle parameters in response to inoculation with different *Pto* DC3000 densities. **C, F, I)** Particle parameters in response to inoculation with different *Pto* DC3000 and co-treatment with flg22. Each dot represents value of independent samples for size and ζ-potential it represents median. 3-12 independent samples were used for each experiment. **J, K)** The profile of ζ-potential for each particle collected from apoplastic fluids of plants treated as indicated and gradient enriched EVs. Control = 0.2 mM EDTA; flg22 = 100 nM; n.t. = not treated; *Pto* DC3000 $OD_{600}$ = 0.0006. Each treatment was 3 days long. The dots represent the mean across the ζ-potential values from independent samples: n = 8 (control); n = 10 (*Pto* DC3000); n = 6 (flg22); n = 4 (non-treatment).

Figure S4

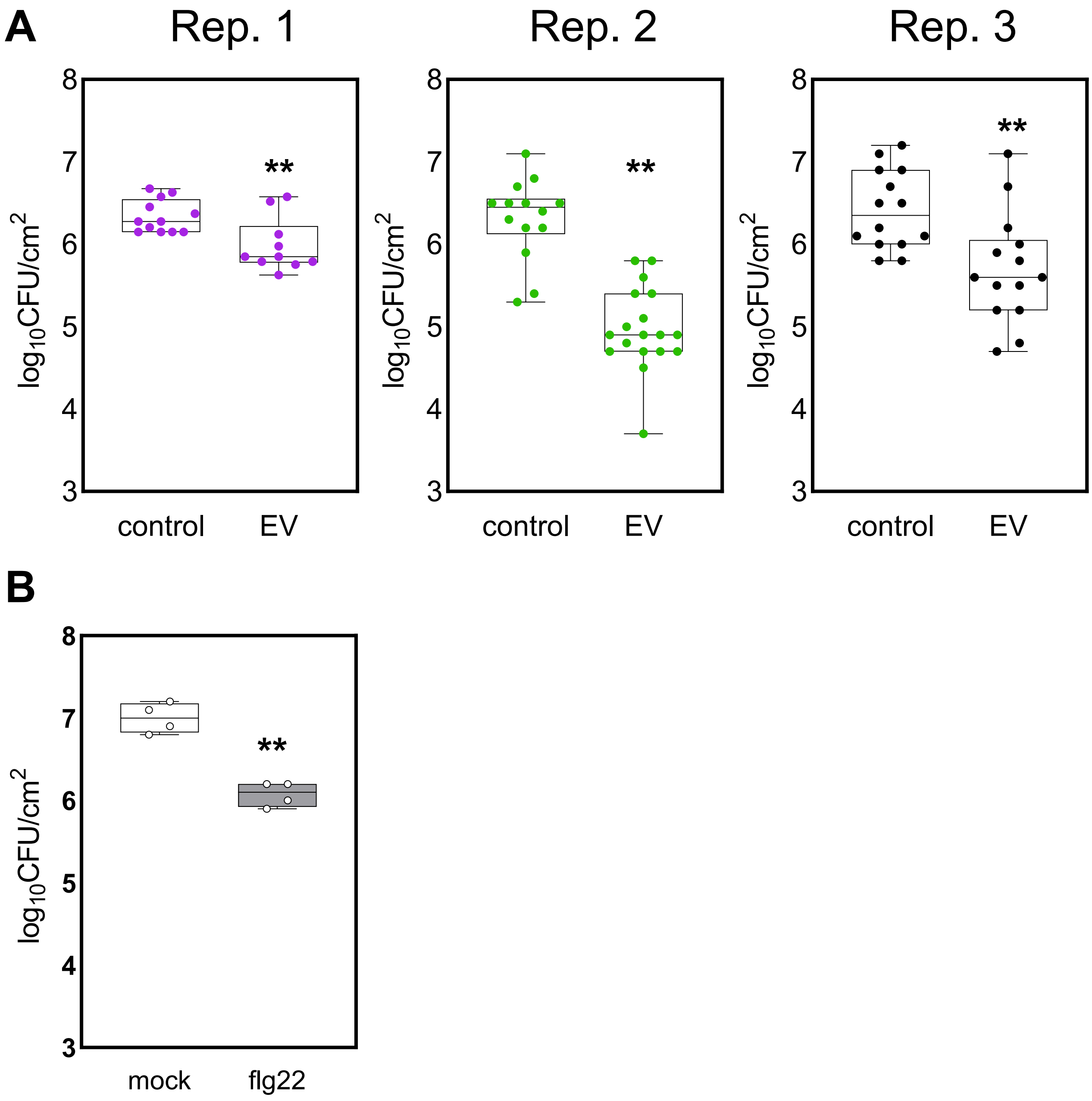


**Figure S4. Pre-treatment with *Pto* DC3000 EVs induces resistance against subsequent *Pto* DC3000 infection. A)** Three individual biological repeats of *Pto* DC3000 growth (CFU) after infection into leaves of *A. thaliana* without and with EV pre-treatment at 3 dpi (control = 0.02 mM EDTA). Each biological repeat consists of 12 independent samples. **B)** *Pto* DC3000 growth (CFU) after infection (3 dpi) into leaves of *A. thaliana* without and with 100 nM flg22 1 day pre-treatment (mock = 10 mM $MgCl_2$) n = 4. Asterisks represent the statistical difference between the treated and control samples (two tailed Student t-test $p < 0.01$).

Figure S5

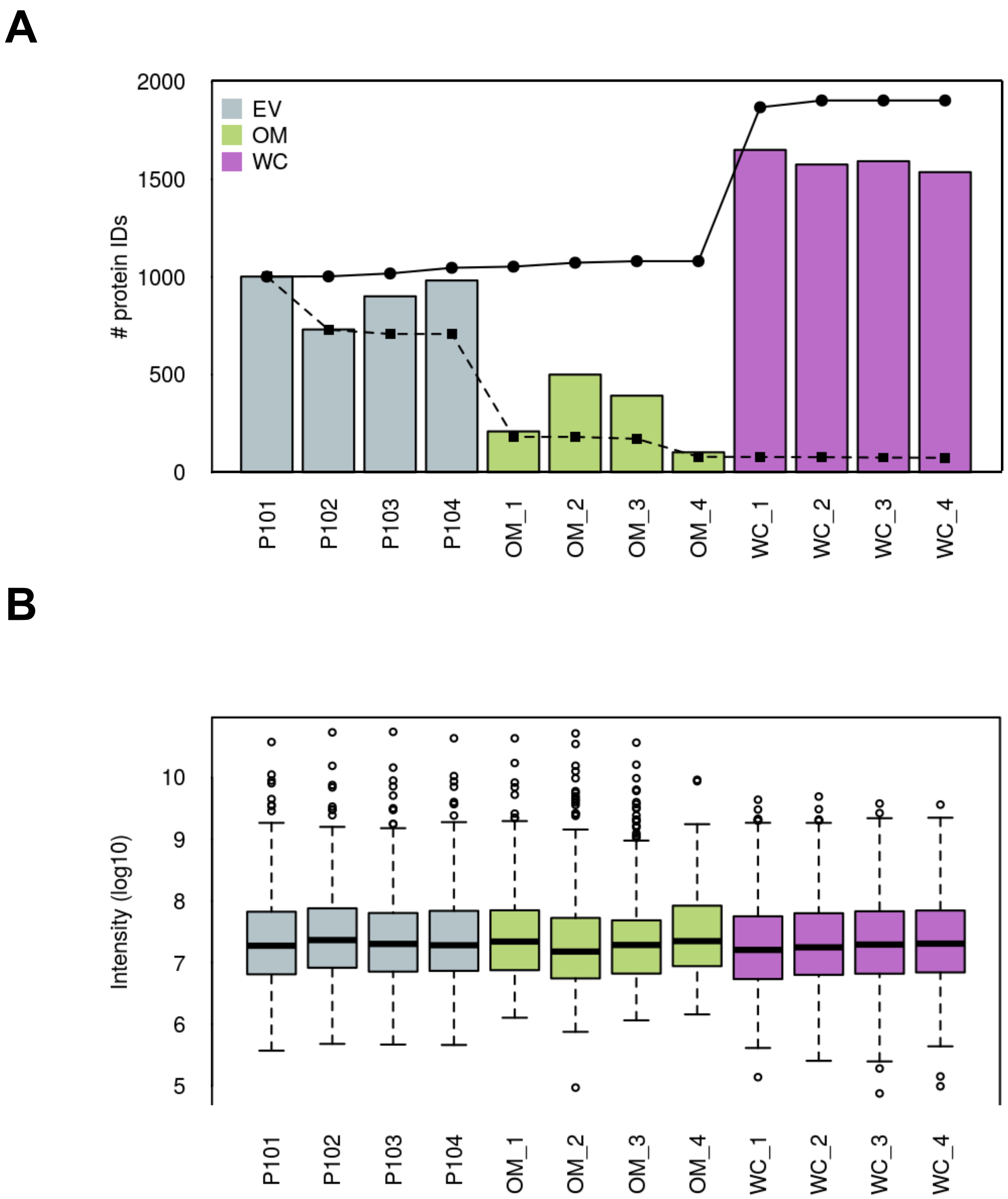


**Figure S5. Characteristics of the proteomic analysis. A)** Barplot shows the number of identified proteins in each replicate. The solid line indicates the cumulative protein IDs and dashed line shows the shared protein IDs. **B)** Boxplot shows a comparable distribution of protein intensities from each replicate.

Figure S6

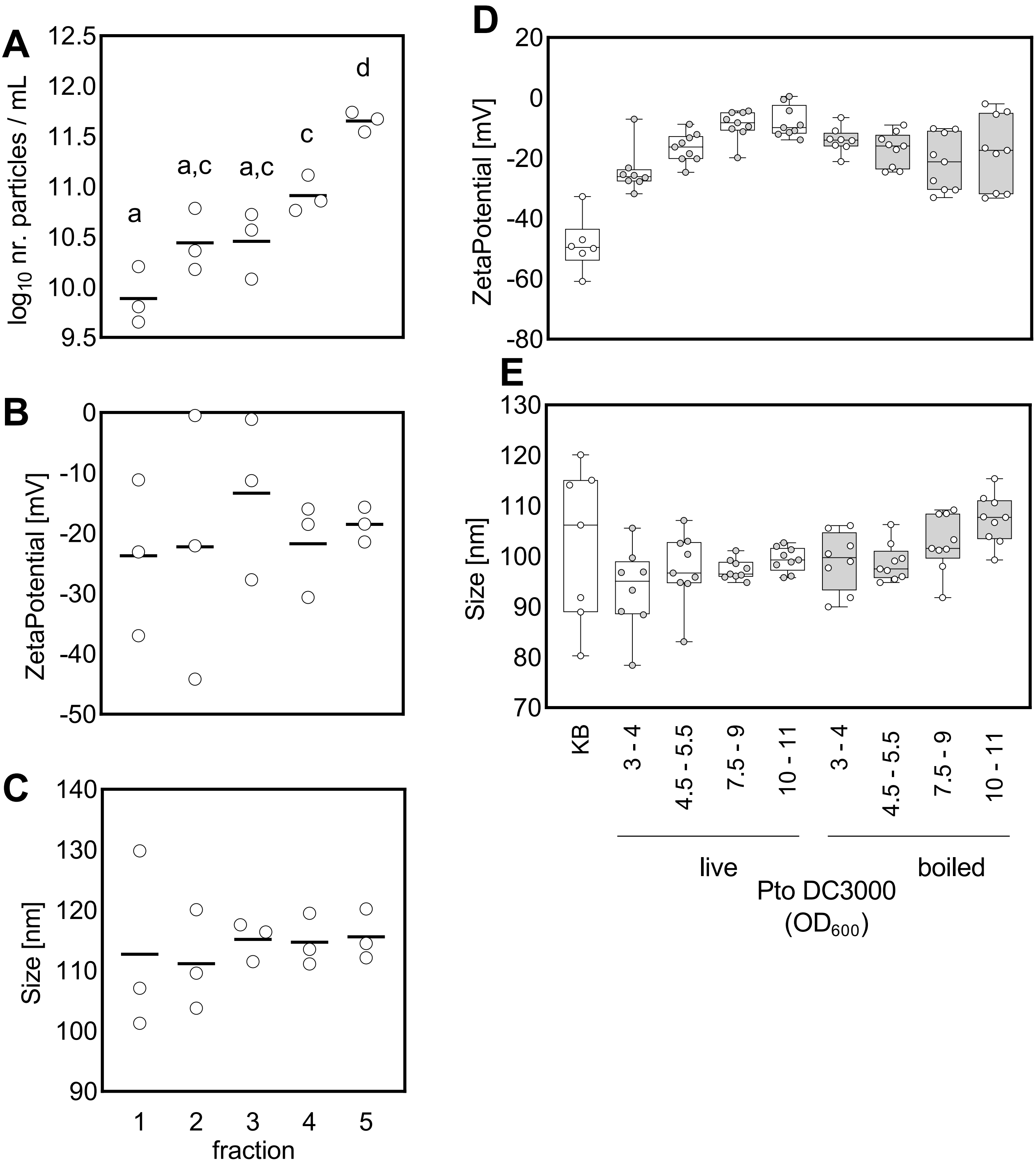


**Figure S6. Biophysical parameters of *Pto* DC3000 EVs across isolation methods. A-C)** NTA measurements of particle concentration (A), ζ-potential (B) and size (C) of *Pto* DC3000 EVs collected from each step of gradient enrichment. **D-E)** NTA analysis of particle ζ-potential (D) and size (E) of *Pto* DC3000 EVs from fluid samples before (live) and after boiling. Each dot represents an independent sample.
